# The locomotor and predatory habits of unenlagiines (Theropoda, Paraves): inferences based on morphometric studies and comparisons with Laurasian dromaeosaurids

**DOI:** 10.1101/553891

**Authors:** Federico A. Gianechini, Marcos D. Ercoli, Ignacio Díaz-Martínez

## Abstract

Unenlagiinae is mostly recognized as a subclade of dromaeosaurids. They have the modified pedal digit II that characterize all dromeosaurids, which is typically related to predation. However, derived Laurasian dromaeosaurids (eudromaeosaurs) differ from unenlagiines in having a shorter metatarsus and pedal phalanx II-1, and more ginglymoid articular surfaces in metatarsals and pedal phalanges. Further, unenlagiines have a subarctometatarsal condition, which could have increased the mechanical efficiency during locomotion. All these discrepancies possibly reflect different locomotor and predatory habits. To evaluate this we conducted morphometric analyses and comparisons of qualitative morphological aspects. The former consisted in two phylogenetic principal component analyses, one of them based on lengths of femur, tibia and metatarsus, and width of metatarsus, and the other based on lengths of pedal phalanges. The data sampling covered several coelurosaurian and non-coelurosaurian taxa. The first analysis showed the unenlagiines close to taxa with long tibiae and long and slender metatarsi, which are features considered to provide high cursorial capacities. Instead, eudromaeosaurs are close to taxa with shorter tibiae and shorter and wider metatarsi, which can be considered with low cursorial capacities. The second analysis showed that eudromaeosaurs and unenlagiines have similar phalangeal proportions. Moreover, they share the elongation of distal phalanges, which is a feature related to the capacity of grasping. The shorter and wider metatarsus, more ginglymoid articular surfaces and a shorter pedal phalanx II-2 of eudromaeosaurs possibly allowed them to exert a greater gripping strength. Thus, they had the potential of hunting large prey. Instead, the longer and slender subarctometatarsus, lesser ginglymoid articular surfaces and a longer pedal phalanx II-2 of unenlagiines possibly gave to them greater cursorial capacities and the ability to hunt smaller and elusive prey on the ground. Thus, the different morphological evolutionary paths of dromaeosaurids lineages seem to indicate different locomotor and predatory specializations.

## Introduction

Unenlagiinae is a clade of Gondwanan paravians first recognized by Bonaparte [1] and which have been generally considered as a subfamily of dromaeosaurids from the phylogenetic analysis made by Makovicky et al. [2]. However, more recently other studies have challenged the dromaeosaurian affinities of unenlagiines and instead have proposed an alternative phylogenetic hypothesis in which these theropods are located within the stem of Avialae [3–4]. Despite this and beyond the discussion about the relationships of unenlagiines, there are many shared morphologic features between unenlagiines and dromaeosaurids. One of these shared traits is the presence of a modified pedal digit II, with a hyperextensible phalanx II-2 and a hypertrophied sickle-shaped claw. The peculiar form of this digit has led many researchers to make multiple interpretations about its possible function (e.g., [5–9]), although they all agree that it was involved in food obtaining, mainly through the submission and/or causing the death of the prey. Nevertheless, these functional interpretations are based mainly on the anatomy of derived Laurasian taxa (i.e., Dromaeosaurinae + Velociraptorinae or Eudromaeosauria following some authors, e.g., [10–11]), such as *Deinonychus*, *Velociraptor*, *Saurornitholestes*, *Achillobator* and *Dromaeosaurus*, in which the phalanges are markedly modified with respect to the plesiomorphic theropod morphology. Regarding the digit II of unenlagiines, it is similarly modified, although there are some anatomic differences with the digit II of eudromaeosaurs.

Moreover, the anatomical differences between unenlagiines and eudromaeosaurs are not limited to those in this pedal digit, but also in other parts of the hindlimb. Mainly, the metatarsus differs between the two groups, since in unenlagiines is generally observed a subarctometatarsal condition, as in microraptorine dromaeosaurids and some basal troodontids, whereas in eudromaeosaurs the metatarsus is more robust and it has a structure more similar to the plesiomorphic condition in theropods. In the subarctometatarsal condition the metapodium has a similar morphology to the arctometatarsus, a type of metatarsal morphology observed in some theropod groups, such as tyrannosaurids, ornithomimids, and alvarezsaurids. White [12] pointed out the way in which both morphologies differ, indicating that in the subarctometatarsus the proximal end of the metatarsal III, although constrained, is equally visible in anterior and plantar views (completely constrained proximally in the arctometatarsus and not visible); and in plantar view the third metatarsal is visible through the entire length of the metatarsus excluding metatarsals II and IV from buttressing. Several functional hypotheses have been raised regarding the arctometatarsus, most of them linked with an increasing of the mechanical efficiency during locomotion [12–17]. The subarctometatarsal condition could have related also to enhance the locomotor efficiency, and some authors consider it as transitional between the plesiomorphic morphology and the arctometatarsal condition [12].

In unenlagiines and eudromaeosaurs the hindlimb, especially the autopodium, is implied both in locomotor and feeding functions, so beyond the phylogenetic relationships between both groups, the morphological differences possibly reflect different locomotor and predatory habits. Based on the previous ideas about the functional implications of the subarctometatarsal and the arctometatarsal condition, likely unenlagiines had locomotor capacities not present in eudromaeosaurs. These hypotheses have already been mentioned by previous authors (e.g., [9]), although not evaluated in a quantitative form, at least not for unenlagiines. The goal of the present contribution is to perform an analysis including taxa of unenlagiines and eudromaeosaurs and to assess, in a quantitative mode, the morphological differences between both groups. Additionally, exhaustive morphological comparisons are performed in order to arrive to a conclusion about the possibly dissimilar functions.

## Materials and methods

In order to evaluate quantitatively how the unenlagiines and eudromaeosaurs differ morphologically was performed a morphometric analysis, employing a set of lineal measurements of the hindlimb bones of several theropod taxa. A diverse sample of theropod clades was considered, including extant birds, with the aim of covering a wide spectrum of morphologies, proportions and sizes of the elements of the hindlimb. So, the sample includes measurements of *Herrerasaurus*, non Tetanurae neotheropods, basal tetanurans, and representatives of most coelurosaur clades including Mesozoic avialans. It was considered also data from more recent although extinct groups of birds, i.e., Dinornithiformes, and from extant taxa, of which the locomotor habit, mode of feeding and capacities of the foot like ‘grasping’ are known. Extant taxa of birds considered include mainly those ground-dwellers with cursorial locomotor habits, raptorial birds with different hunting modes and ‘grasping’ capacities, and perching birds with more arboreal habits, such as passeriforms, also with ‘grasping’ capacities (S1 Appendix). The measurements considered included proximodistal lengths of the femur (FL), tibiotarsus (TL), metatarsus (MtL), and non-ungual pedal phalanges, and the lateromedial width of the metatarsus at midshaft (ML). Regarding to MtL, the measures were taken for the longest element, typically the metatarsal III. For modern birds was considered the length of the tarsometatarsus, due to the complete fusion of the distal tarsals and metatarsals. The dimension ML refers to the lateromedial diameter of the articulated MT II, III and IV at midshaft of these bones.

Most of values of these measurements were obtained from previously published datasets, especially from [15] and also from other authors (see supplementary information), whereas others were obtained directly from materials deposited in different collections. For many taxa with published measurements the dimension ML was not considered by the authors, so in these cases ML was calculated from the published photographs of the specimens. For each taxon is specified the specimen from which the measurements were taken, except some not indicated by the author who published the data. In the case of taxa for which there are published measurements of several specimens, it has been decided to consider the data of only one of them, specifically the larger one, in order to avoid data of juvenile forms. Jointly, those specimens that were as complete as possible were taken into account, i.e., those with all the bones of the hindlimb preserved completely, in order to obtain the data of all the measurements. In some cases, estimated measurements have been taken of bones that have a small part not preserved, so even if it is estimated it is quite approximate to the real one. Additionally, measurements were obtained directly from materials housed in repositories of Argentina, including one specimen of the alvarezsauroid *Alnashetri cerropoliciensis* (MPCA), 17 specimens of many taxa of extant birds (MACN), and one specimen of *Struthio camelus* (CFA-OR). These are specified in the S1 Appendix.

Regarding lengths of pedal phalanges they were not taking into account the lengths of unguals, because there is no a consensus on how to measure this length, since some authors measure it in a straight line from the proximal end to the distal end of the phalanx, while others measure only the external curvature. So, published lengths of pedal unguals of theropods are not taken with the same criteria. Neither was considered the lengths of the phalanges of digit I, because in taxa of some clades included in the analysis, i.e., ornithomimids, this digit is reduced and completely absent.

From this measurements phylogenetic principal component analyses (Phylogenetic PCA; see [18–19]) instead traditional PCA were performed. The phylogenetic principal component analyses allow the reduction of original variables to principal components correcting the non-independence among the former due the phylogenetic relationships between species. In this way, in a phylogenetic PCA the samples are not considered as independent datapoints, an assumption of the traditional PCA and frequently violated due the phylogenetic relationships between samples [18].

Given that the purpose of these analyses was the study of shape changes between species that cover a wide diversity of sizes, the phylogenetic PCA were constructed from, size-standardized, Mosimann variables [20], instead original ones. Each Mosimann variables were obtained as the ratio between the original variable and the geometric mean of all variables considered for the corresponding phylogenetic PCA.

From the complete dataset two phylogenetic principal component analyses were performed. One of them includes the long bones of the hindlimb measurements, i.e., FL, TL, MtL and ML, and the other one includes the lengths of the non-ungual pedal phalanges. In relation to the available data (S1 Appendix), the first PCA included 74 taxa, whereas the second one 32 taxa. This analysis design implying different taxonomic representatives in each principal component analysis (in relation to the available data and inability to perform these analyses with missing data), but allowed the maximization of the number of morphologies and taxa considered in each analysis.

After computed to Phylogenetic PCA, the phylogenetic relationships between species were projected into bivariate plots of morphospaces, constructing phylomorphospaces [19]. To evaluate the phylogenetic signal on each phylogenetic principal component, the K statistic proposed by Blomberg et al. [21] where calculated for each axis. The K statistic provides a measure of the strength of phylogenetic signal data. The values smaller than one indicate a lack of phylogenetic signal or strong adaptative processes, values near 1 are expected if the character evolved following the phylogenetic relationships, under a Brownian motion model, and values greater than one show that phylogenetically closer taxa are more similar than expected, and eventually stasis processes [21–22].

Additionally, the size-effect on each axis of the morphospaces were calculated using phylogenetic generalized least squares (PGLS) regressions [23], considered the geometric mean as the independent variables. A PGLS regression allows the incorporation of the phylogenetic structure of samples as the error term of the regression equations, and then considering the biases caused by phylogeny in the calculation of the relationship between the analyzed variables.

All these analyses were carried out using the software R 3.5.0 [24] and using the PHYTOOLS [19], APE [25], and PICANTE [26] libraries.

For the Phylogenetic PCA and the PGLS, both for the analysis based on long bones measurements and that based on lengths of the phalanges, were used composited phylogenies which synthetized the relationships between taxa included in the study. These were based on previously published phylogenies of different theropod clades [27–42].

The morphological differences between unenlagiines and other dromaeosaurids also were evaluated through qualitative comparisons of the hindlimb bones, especially of the matatarsals and pedal phalanges. The morphology of dromaeosaurid taxa was observed directly from the holotypes of *Deinonychus* (YPM 5205), *Bambiraptor* (AMNH FR 30556), and *Dromaeosaurus* (AMNH FR 5356), and from the literature (e.g., [5–6, 10, 35, 40, 43–52]). The observations of the unenlagiines were made on the holotypes and referred materials of *Buitreraptor* (MPCA 245, MPCA 238, MPCA 478, and MPCN-PV-598), *Neuquenraptor* (MCF PVPH 77), *Austroraptor* (MML 195 and MML 220), and a cast of the holotype of *Rahonavis* (FMNH PR 2830). Additional comparisons with other theropod taxa were made using the literature and in the case of extant birds also using the materials above mentioned.

Curvature angles of unguals of unenlagiines and Laurasian dromaeosaurids were measured using the methodology applied by Fowler et al. [53], which in turn is based on that of Pike and Maitland [54]. Both the external and inner curvature angles of the unguals are measured with this methodology, i.e., from the dorsal and ventral borders respectively, obtaining the angle between the base and the tip of the claw. However, as this methodology was used to measure ungual curvatures of extant taxa of birds with soft tissue on digits some modifications were made. For extant birds the base of the claw is considered the point where the keratinous sheath emerges from the skin of the digit, although in fossil unguals lacking the sheath and soft tissue cannot be considered the same base of the claw to the measurement of the curvature angles. So, we take the proximodorsal tip of the ungual bone as the dorsal base to measure the external curvature angle, and the tip of the flexor tubercle as the ventral base (S2 Fig). However, the flexor tubercle shows two ventral tips in unguals of the analyzed theropods, both separated by an extension of the side groove of the claw, so the anterior end was taken as the base to measure the angle of the inner curvature. The angles were taken from photographs of the ungual phalanges using the measure tool in Adobe Photoshop. For incomplete materials which have not preserved the distal or the proximoventral ends these were reconstructed, although in these cases was indicated that the angle values are estimated.

### Institutional abbreviations

AMNH FR, American Museum of Natural History, New York, NY, USA. CFA-OR, Fundación de Historia Natural “Félix de Azara”, Ciudad Autónoma de Buenos Aires, Argentina. FMNH PR, Field Museum of Natural History, Chicago, IL, USA. MACN, Museo Argentino de Ciencias Naturales “Bernardino Rivadavia”, Ciudad Autónoma de Buenos Aires, Argentina. MCF PVPH, Museo “Carmen Funes”, Plaza Huincul, Neuquén, Argentina. MML, Museo Municipal de Lamarque, Lamarque, Río Negro, Argentina. MPCA, Museo Provincial “Carlos Ameghino”, Cipolletti, Río Negro, Argentina. MPCN-PV, Museo Patagónico de Ciencias Naturales, General Roca, Río Negro, Argentina. MUCPv, Museo de Geología y Paleontología de la Universidad Nacional del Comahue, Neuquén, Argentina. YPM, Yale Peabody Museum, New Haven, CT, USA.

## Results

### Description of the PPCA based on hindlimb long bones measurements

In the PPC analysis based on hindlimb long bones (femur, tibia, and metatarsals) measurements, including Mesozoic theropods (MzTer), extant birds, and Dinornithiformes the contributions of the osteological variables to the first principal component (PPC1) represent 57.2% and to the second principal component (PPC2) represent 30.1% of the total variation (Fig 1). The PPC1 summarizes a major contribution of tibia and metatarsus lengths (negatively correlated with the PPC1) and the mediolateral width of metatarsus at midshaft (ML; positively correlated). High negative PPC1 scores depicted taxa with elongated and slender metatarsi and elongated tibiae, whereas less negative and positive PPC1 scores depicted taxa with shorter and wider metatarsi and shorter tibiae. The PPC2 summarizes a major contribution of femur length (positively correlated) and minor contributions of metatarsus length and ML (both variables negatively correlated). High positive PPC2 scores depicted taxa mainly with elongated femora and slightly short and slightly slender metatarsi, whereas negative scores depicted taxa mainly with shorter femora and slightly longer and slightly wider metatarsi.

**Fig 1.**
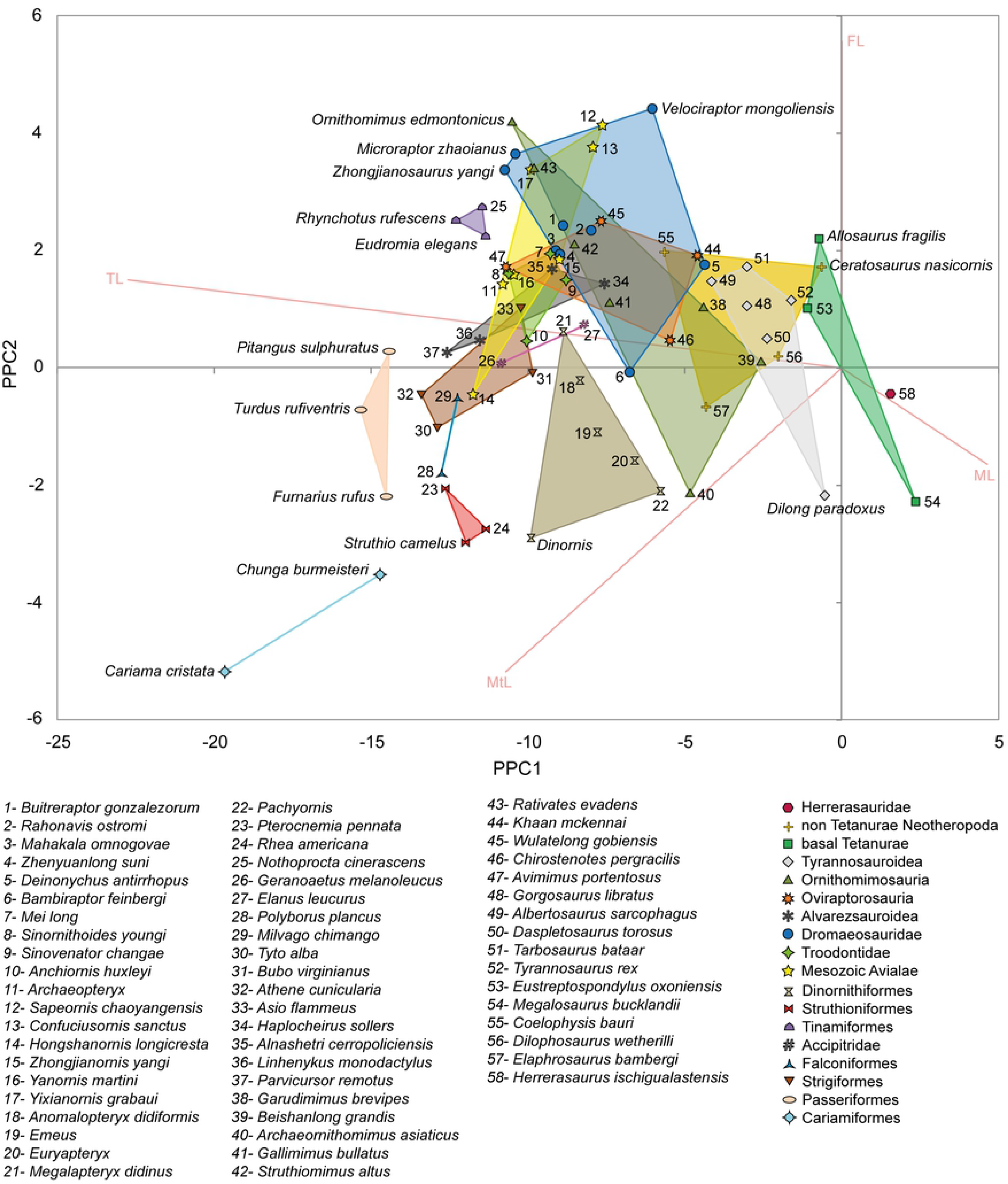
Morphospace obtained from the phylogenetic PCA based on the measurements of the hindlimb long bones.

Extant birds and Dinornithiformes are partially segregated from the MzTer, toward negative scores of PPC1 and PPC2 (Fig 1). This is mainly because these groups of birds have comparatively longer and more slender metatarsi, longer tibiae, and shorter femora in comparison with the MzTer. More specifically, those taxa showing the longest and more slender metatarsi include terrestrial flightless or sparingly flying birds, i.e., the Cariamiformes (*Cariama* and *Chunga*) and the Struthioniformes, as well as some Passeriformes such as *Furnarius*. In fact, the Cariamiformes stand out by having extremely elongated metatarsi and tibiae, with a metatarsus longer than 1.5 times the femur length or even more than twice longer than the femur (as in *Cariama*) and a tibia longer than twice the femur length. The remaining terrestrial birds, i.e., the Tinamiformes and Dinornithiformes, show comparatively shorter metatarsi and tibiae. The Tinamiformes are located on negative PPC1 and positive PPC2 scores, closely to the arcto and subarctometatarsalian MzTer with elongated metatarsi and tibiae. With respect to the Dinornithiformes, they have comparatively wider metatarsi than the Tinamiformes.

Some extant raptor birds, such as the accipitrids (*Elanus* and *Geranoaetus*), some Strigiformes (*Asio* and *Bubo*), and some Dinornithiformes (*Megalapteryx*) are on low negative and positive PPC2 scores, closer to the MzTer with the longest and more slender metatarsi and the longest tibiae. These taxa show short and wide metatarsi, when are compared with the remaining extant birds (except the Tinamiformes).

Regarding to the MzTer, on high negative PPC1 scores are located the alvarezsauroids, derived ornithomimids, some oviraptorosaurs, basal avialans, troodontids, microraptorines, and unenlagiines, all of them with markedly elongated hindlimbs (more elongated and slender metatarsi and longer tibiae in comparison with the remaining MzTer). Moreover, many of these taxa are characterized by an arctometatarsal or subarctometatarsal condition.

The MzTer with the longest and the more slender metatarsi and longest tibiae are located on the highest negative PPC1 scores and among the lowest positive and some negative PPC2 scores. These taxa include the derived alvarezsaurids *Parvicursor* and *Linhenykus*, both with a very slender and markedly elongated arctometatarsalian metatarsus, which significantly surpass the femur length. Also is included in this part of the morphospace the basal avialan *Hongshanornis*, which although does not have an arctometatarsalian condition present a notably elongated and slender metatarsus which equals the femur length, locating on negative PPC2 scores (differing from other basal avialans).

Regarding unenlagiines, *Buitreraptor* is closer to *Mahakala*, *Zhongjianornis*, *Zhenyuanlong*, *Struthiomimus*, *Mei*, *Alnashetri*, and *Sinovenator* (Fig 1). These taxa show a long metatarsus, although slightly shorter and wider than in the MzTer above mentioned, so they are located on less negative PPC1 scores and more positive PPC2 scores. *Rahonavis* is closer to the oviraptorosaur *Wulatelong* than to *Buitreraptor* and presents less negative PPC1 scores and slightly lower positive PPC2 scores than *Buitreraptor*. This separation is because *Rahonavis* has a slightly shorter and wider metatarsus than *Buitreraptor* and the other taxa closer to it.

*Deinonychus* and *Velociraptor* segregate and locate on less negative PPC1 scores than other dromaeosaurids, including *Buitreraptor*, since they have markedly shorter and wider metatarsi and shorter tibiae. In fact, *Deinonychus* is closer to tyrannosaurids than to other dromaeosaurids. *Velociraptor* is located on much more positive PPC2 scores, because it has an even shorter metatarsus and tibia comparatively with the femur. *Khaan* is an oviraptorid with hindlimb bones proportions similar to *Deinonychus*.

The large-sized tyrannosaurids have short and wide metatarsi and short tibiae, although an arctometatarsalian condition. Many non-arctometatarsalian taxa characterized by relatively elongated although moderately wide metatarsi and moderately elongated tibiae, are located on low negative PPC1 scores and on low positive and negative PPC2 scores. Among these taxa are included basal ornithomimosaurs, the oviraptorosaur *Chirostenotes*, the ceratosaur *Elaphrosaurus*, and the dromaeosaurid *Bambiraptor*. It is noteworthy that *Bambiraptor* is separated from the other derived Laurasian dromaeosaurids, mainly due it shows a comparatively longer metatarsus.

The basal tetanurans, ceratosaurs, coelophysoids, and *Herrerasaurus* are located on the lowest negative and positive PPC1 scores. These taxa have a foot with plesiomorphic morphology showing the shortest and widest metatarsi and shortest tibiae among the theropods included in the analysis. The tyrannosauroid *Dilong* has a longer metatarsus with a more derived morphology, although it remains closer and is grouped with the mentioned taxa due its relatively wide metatarsus.

### Influence of phylogeny in the distribution of taxa along the morphospace

The K of Blomberg values indicate that the taxa distribution along the PPC1 is strongly influenced by the phylogenetic relationships of major clades (K=2.714) whereas PPC2 is less influenced by deep phylogenetic relationship, and related to the influence of the phylogenetic structure of terminals and more inclusive clade (K= 0.262) (S3 Table). Thus, the segregation and relatively scarcely overlapped distribution of these major clades along the PPC1 can be related to the high K value of this axis, while the low K value of PPC2 indicates that there exist many convergences to extreme values in different terminal and less inclusive clades. Observing the phylogenetic relationships plotted on the morphospace (i.e., phylomorphospace; Fig 2) there exist a main separation trend of birds (including extant taxa and Dinornithiformes) toward negative values of the PPC1 and MzTer in less negative and positive values of PPC1. This separation is because birds have generally a longer and slender metatarsus and a longer tibia than MzTer. In addition, more derived taxa of some MzTer clades generally trend to locate on more negative values of PPC1 (as can be observed in tyrannosauroids, ornithomimosaurs, and alvarezsaurs), while most primitive taxa considered locate on the extreme positive values of PPC1, in relation with their plesiomorphic metatarsal morphology (Fig 2).

**Fig 2.**
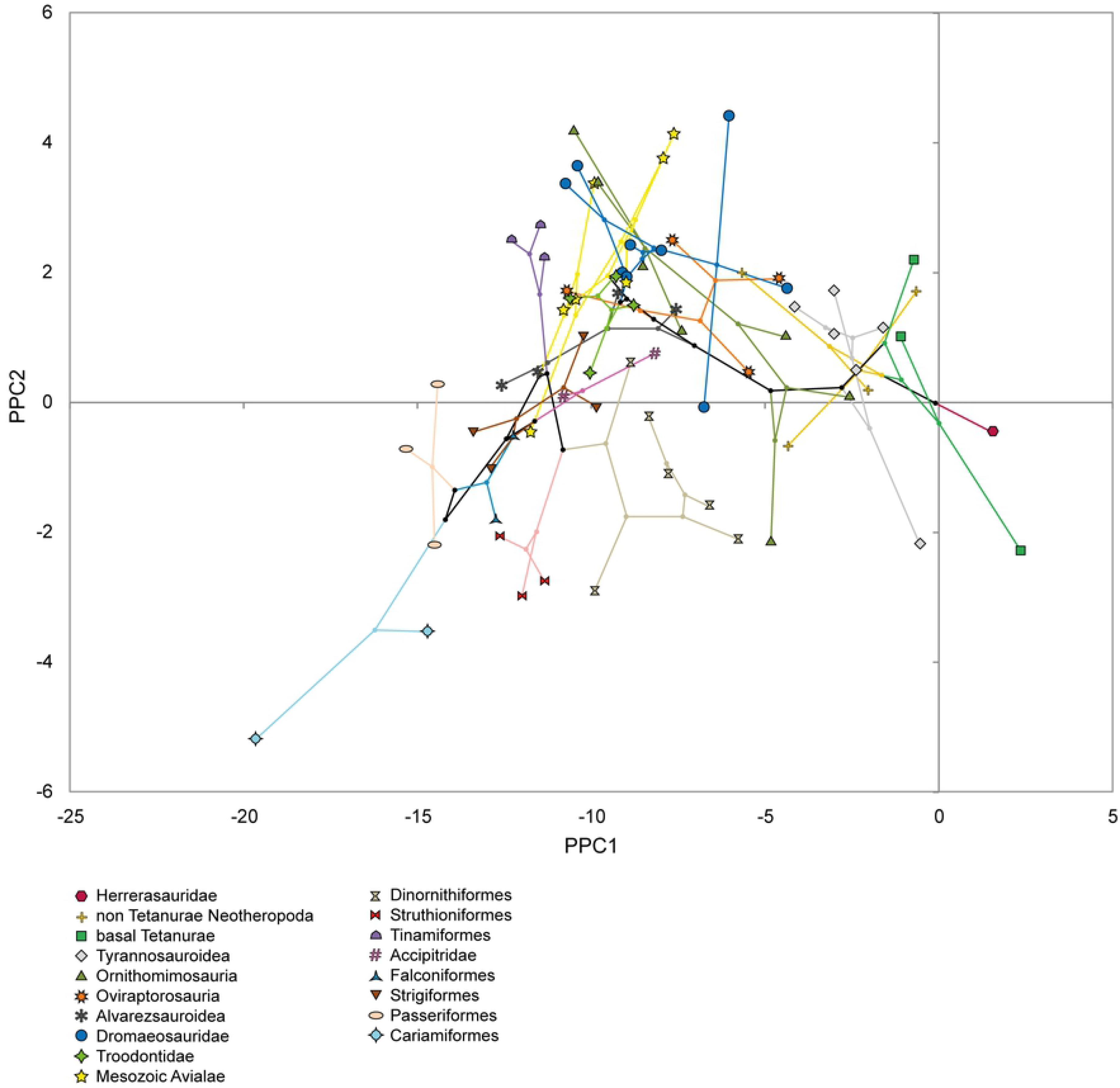
Phylomorphospace obtained from the phylogenetic PCA based on the measurements of the hindlimb long bones.

As was stated above, PPC2 summarizes morphological similarities between minor clades or terminals. Although PPC2 is less influenced by the structure of phylogenetic relationships of major clades, the positive correlation of this component with the femoral length can partially explain the division between MzTer and extant birds and Dinornithiformes, because in the latter there is a general trend to a significant shortening of the femur in comparison with MzTer, reason why they are mostly on negative values of the PPC2. The exception is the Tinamiformes, which are on positive values of PPC2, significantly separated from the remaining modern birds.

Regarding the distribution of MzTer along the PPC2 some trends are observed. Thus, in tyrannosauroids and ornithomimosaurs there is a marked separation between basal and derived taxa, because basal taxa are located on negative values (with short femora) and derived taxa tend to more positive values (with longer femora). Derived ornithomimosaurs and alvarezsauroids are characterized by an elongated tibia and metatarsus, although in the former the trend was a comparatively more marked elongation of the femur whereas in alvarezsauroids the trend was to shorten the femur in derived forms. In oviraptorosaurs the direction of the morphological changes was not as clear as in the coelurosaur groups mentioned, possibly because the small taxon sample is not adequate to show a clearer trend. Troodontids show a distribution of taxa along PPC2 similar to tyrannosauroids and ornithomimosaurs, since basal taxa are on low positive values of the axis whereas more derived taxa are on more positive values and hence they present a longer femur.

About the distribution of dromaeosaurids along the phylomorphospace (Fig 2) is observed an opposite tendency in comparison with other groups mentioned, since more basal taxa, such as *Mahakala* and the unenlagiines *Buitreraptor* and *Rahonavis*, are located on more negative values of PPC1, whereas more derived taxa, i.e., *Deinonychus*, *Velociraptor*, and *Bambiraptor*, are on less negative values of PPC1. In this way, the basal taxa present a longer metatarsus and tibia than derived forms. Regarding the location of taxa along PPC2, dromaeosaurids not show a clear trend, in contrast to the clades already explained. Basal taxa are located on similar values of PPC2, whereas microraptorines (at least those considered in this analysis) are more widely distributed. Thus, some microraptorines (i.e. *Microraptor* and *Zhongjianosaurus*) are on high positive values of PPC2, whereas others (i.e. *Zhenyuanlong*) are on similar PPC2 values than basal dromaeosaurids, with a shorter femur. Moreover, *Microraptor* and *Zhongjianosaurus* converge in the morphospace with derived ornithomimids with long femora. The derived dromaeosaurids are also widely distributed, being *Velociraptor* on high positive values of PPC2, close to some basal avialans, *Bambiraptor* on negative values, close to taxa with a shorter femur and longer metatarsus, and *Deinonychus* on an intermediate location near derived tyrannosaurids. The location of *Velociraptor* can be explained possibly by its comparatively longer femur with respect to the other derived dromaeosaurids here analyzed.

### Influence of size in the distribution of taxa along the morphospace

The PGLS regressions indicates that the PPC1 in the analysis based on long bones dimensions is significantly influenced by size (F = 7.318; p-value = 0.009). The MzTer taxa with the largest body sizes are located to the right side of the morphospace, on less negative and some positive values of the PPC1. Furthermore, Dinornithiformes, the larger modern birds considered in the analysis, are located to the right of the morphospace occupied by birds. These large-sized taxa are characterized by a comparatively short and wide metatarsus, as was explained above. By other side, smaller taxa with slender and longer hindlimbs are situated at the left of the morphospace, whether in the case of MzTer or modern birds. Conversely, PPC2 (F = 2.162; p-value = 0.146) and PPC3 (F = 3.260; p-value = 0.075) in not significantly influenced by size, and it result agrees with the distribution of taxa along the axis.

### Description of the PPCA based on phalanges lengths

In the PPC analysis made from phalanges lengths, the contributions of the variables to the first principal component (PPC1) represent 39.0% and to the second principal component (PPC2) represent 29.1% of the total variation (Fig 3). Because these two axes explain a small percentage of the variation, we also analyzed the third component (10.8% of the total variation).

**Fig 3.**
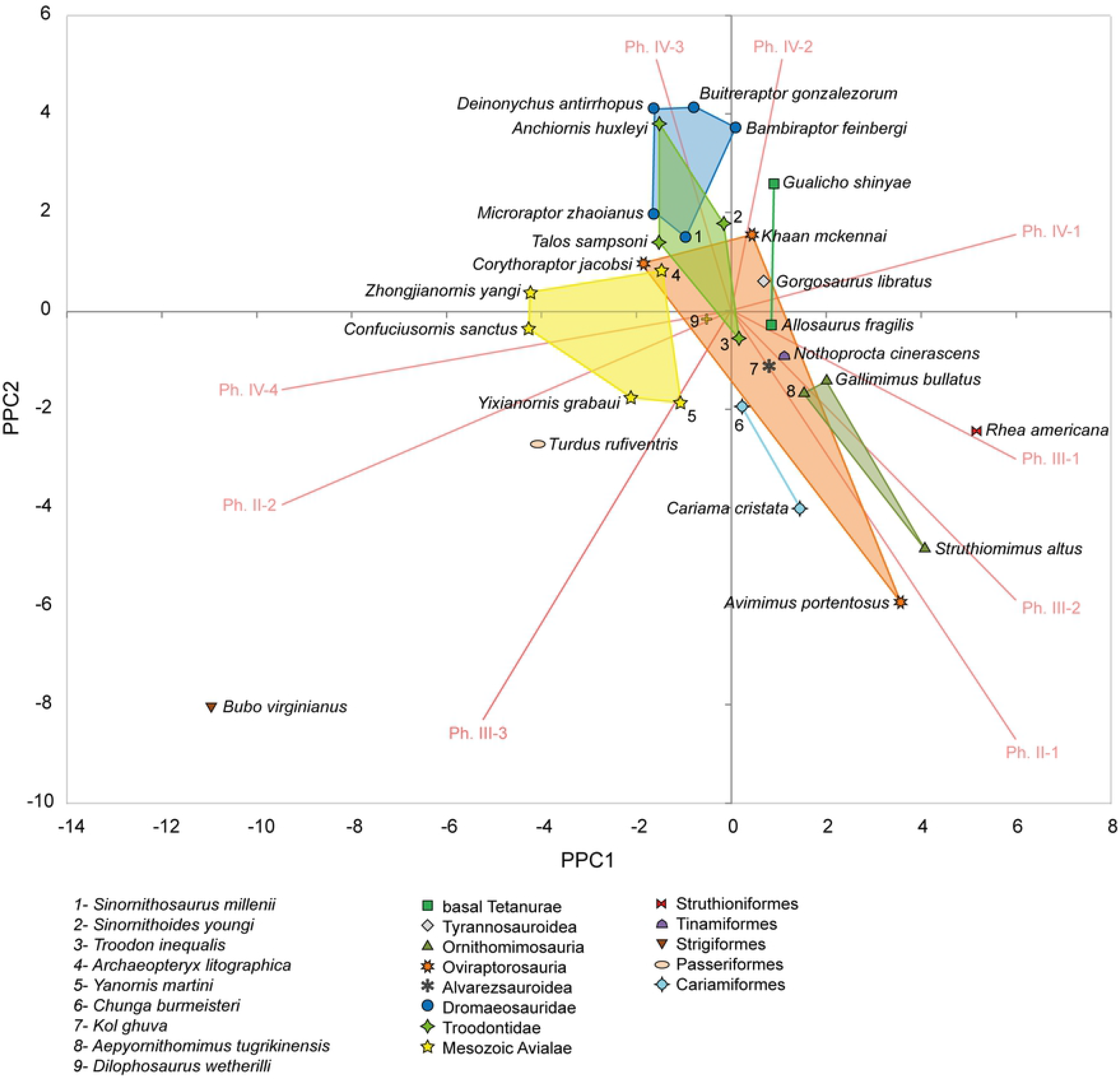
Morphospace obtained from the phylogenetic PCA based on the lengths of the pedal phalanges (PPC1 vs PPC2).

In the graphic of PPC1 vs PPC2 (Fig 3), the PPC1 summarizes a major contribution of the lengths of the proximal phalanges, i.e., Ph. II-1, III-1, IV-1, and III-2 (positively correlated with this component), and the lengths of the distal pre-ungual phalanges, i.e., II-2, III-3, and IV-4 (negatively correlated with this component). In this way high positive PPC1 scores depict taxa with elongated proximal phalanges and high negative PPC1 scores depict taxa with elongated distal phalanges. The PPC2 summarizes major contributions of the lengths of the proximal and middle phalanges of digit IV, i.e., IV-2 and IV-3 (positively correlated with this component), and the lengths of the proximal and distal pre-ungual phalanges of digits II and III (negatively correlated). Thus, high positive PPC2 scores depict taxa with long phalanges IV-2 and IV-3 whereas high negative PPC2 scores depict taxa with long proximal or distal pre-ungual phalanges. Considering both principal components and summarizing the distribution along the morphospace, taxa on high positive PPC1 and high negative PPC2 scores are between those showing more elongated proximal phalanges, whereas those taxa located on negative PPC1 scores have relatively more elongated distal phalanges.

For the graphic of PPC2 vs PPC3 (Fig 4), in addition to those already commented for PPC2, the PPC3 (10.8% of the total variation) summarizes major contributions of the lengths of all the phalanges of digit II (positively correlated with this component), and to a lesser extent it summarizes contributions of the lengths of phalanges of digit III, mainly Ph. III-2 and III-3 (negatively correlated). Thus, high positive PPC3 scores depict taxa with a long digit II whereas high negative PPC2 scores depict taxa with long phalanges III-2 and III-3.

**Fig 4.**
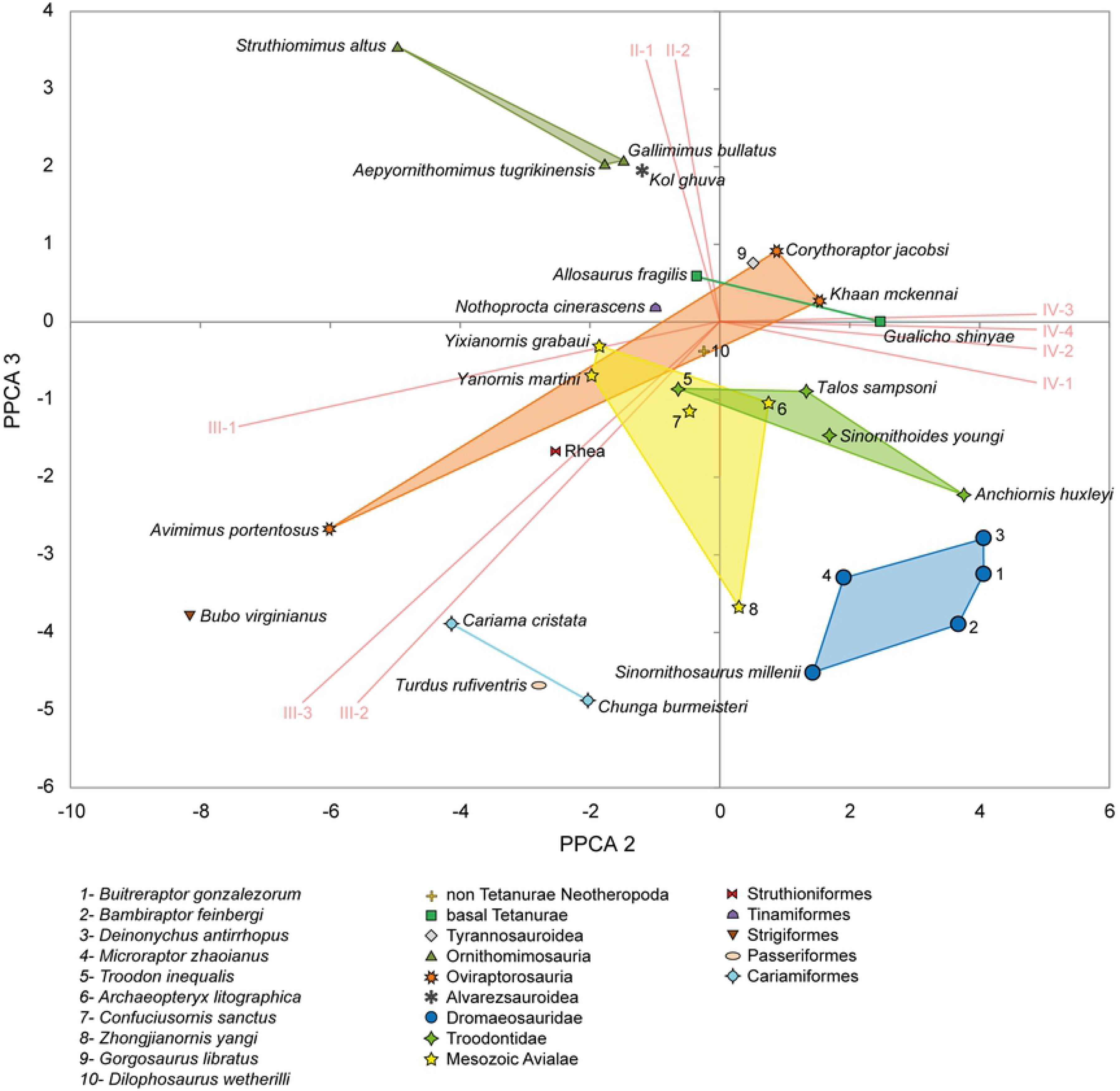
Morphospace obtained from the phylogenetic PCA based on the lengths of the pedal phalanges (PPC2 vs PPC3).

In the morphospace, dromaeosaurids occupy central values of PPC1, high positive of values of PPC2, and high negative PPC3 scores (Figs 3 and 4). All representatives are mainly on negative PPC1 scores, except *Bambiraptor* which is on low positive PPC1 values. *Deinonychus*, *Buitreraptor*, and *Bambiraptor* are located on higher positive PPC2 scores and *Microraptor* and *Sinornithosaurus* on less positive scores of this component. The high positive values of PPC2 of dromaeosaurids are linked to a remarkably elongated digit IV, a feature mainly product of elongation of phalanges IV-2 and IV-3, while the high negative values on PPC3 are also mainly related to the length of phalanges of digit IV, but also influenced by the length of Ph. III-2 and III-3. *Deinonychus* and *Buitreraptor* show a relatively long digit IV in comparison with other dromaeosaurids, although *Deinonychus* is slightly located on more negative PPC1 scores so the position of this taxon is also specifically influenced by the length of phalanx IV-4. *Sinornithosaurus* is the taxon with higher PPC3 values, a position influenced by the elongated phalanges III-3 and II-2. The location of *Microraptor* is due a relatively shorter digit IV in comparison with *Deinonychus*, *Buitreraptor*, and *Bambiraptor*, whereas the length of phalanx IV-4 influenced in its position on more negative PPC1 scores.

Troodontids show a distribution on the morphospace mainly similar to that of dromaeosaurids (Figs 3 and 4), except by *Troodon* which is located on negative PPC2 scores, with relatively shorter digit IV than the others troodontids in the analysis. *Anchiornis* is much close to *Deinonychus* and *Buitreraptor*, a position mainly influenced by a long digit IV. The location of *Sinornithoides* and *Talos*, in less negative values of PPC2, is related to their less elongated digit IV in comparison with *Anchiornis*. In turn, *Talos* is close to *Microraptor* and thus its position is also influenced by the length of phalanx IV-4.

Non-coelurosaurian theropods are dispersed in the morphospace (Figs 3 and 4). *Dilophosaurus* locates on very low negative PPC1, PPC2, and PPC3 scores, showing subtle elongated distal phalanges and a slightly longer digit III. The two basal tetanurans included in the analysis, i.e. *Allosaurus* and *Gualicho*, have similar PPC1 and PPC3 values, although they segregate along PPC2, thus indicating that the difference in length between digit IV and digits II and III is the main factor influencing the separation of these tetanurans. The position of *Gorgosaurus*, which is the only tyrannosaurid included in the analysis, is mainly influenced by relatively long proximal phalanges and especially by those of digit IV.

Oviraptorosaurs show a wide distribution on the morphospace (Figs 3 and 4), since *Corythoraptor* and *Khaan* have a more elongated digit IV whereas *Avimimus* has more elongated proximal phalanges of digits II and III and a comparatively longer digit III.

Ornithomimosaurs are on positive PPC1 and PPC3 scores and on negative PPC2 scores, a location mainly influenced by longer proximal phalanges and a relatively longer digit II. The position of *Struthiomimus* is related to a longer digit II and Ph. III-1 than those of *Gallimimus* and *Aepyornithomimus*.

Mesozoic avialans are on negative PPC1 and PPC3 scores and on positive and negative PPC2 scores (Figs 3 and 4). Basal taxa, i.e., *Archaeopteryx* and *Zhongjianornis*, are on positive PPC2 values, although *Zhongjianornis* highlights due it is located on high negative PPC3 values. The position of *Archaeopteryx* indicates that it has a long digit IV, mainly due elongation of Ph. IV-3 and IV-4. The location of *Zhongjianornis*, *Confuciusornis*, *Yanornis*, and *Yixianornis* is mainly influenced by a greater elongation of distal phalanges of digits II, III, and IV and by a digit III comparatively longer. Specifically, the position of *Zhongjianornis* is biased by the length of digits III-2 and III-3, and secondarily influenced by a long digit IV, whereas the location of *Yanornis* and *Yixianornis* is more influenced by a longer Ph. III-1 and the position of *Confuciusornis* is secondarily influenced by a longer digit IV.

Extant birds are mainly distributed along negative PPC2 and PPC3 scores, although there is observed a dichotomy along the PPC1, because some taxa are on positive scores and others on negative ones (Figs 3 and 4). The position of taxa on positive PPC1, such as *Rhea*, *Nothoprocta*, *Cariama*, and *Chunga* is mainly influenced by long proximal phalanges of digits II, III, and IV. The most notorious bias is observed on *Rhea*, whilst the position of *Cariama* and *Chunga* is also influenced by the length of Ph. III-3. Those taxa on negative PPC1 scores, i.e., *Turdus* and *Bubo* are markedly influenced by the length of distal phalanges of digits II, III, and IV, being the position of *Bubo* the more affected by this trait. Additionally, the position of these two taxa is biased by a comparatively longer digit III. Moreover, this digit is longer in *Bubo* and *Turdus* in comparison with digit III of the other extant birds analyzed.

### Influence of phylogeny in the distribution of taxa along the morphospace

The K of Blomberg values indicate that the taxa distribution along the PPC1, PPC2, and PPC3 is strongly influenced by relationships between terminals and less inclusive clades in the case of PPC1 (K=0.303) and PPC2 (K=0.376), and linked to the many morphological convergences between distant taxa described above (S4 Table). The PPC3 also show a K value lesser than 1 but more closer to 1 (K=0.811), fitting more closely with a stochastic model (i.e., the distribution of taxa follows the phylogenetic relationships but is not particularly strong influenced by deep nodes neither terminal relationships).

For instance, basal taxa included in the analysis, such as basal tetanurans and the basal coelurosaur *Gorgosaurus* are almost indifferently located on similar values of PPC1, although they are separated along PPC2 and PPC3 (Figs 5 and 6). The ornithomimids also are significantly separated mainly along PPC2, although the scarce sample of this group of theropods and recent phylogenetic analyses [55], which show them in a polytomy on the cladogram, difficult to shed light to how phylogeny and the distribution along the mosphospace are related.

**Fig 5.**
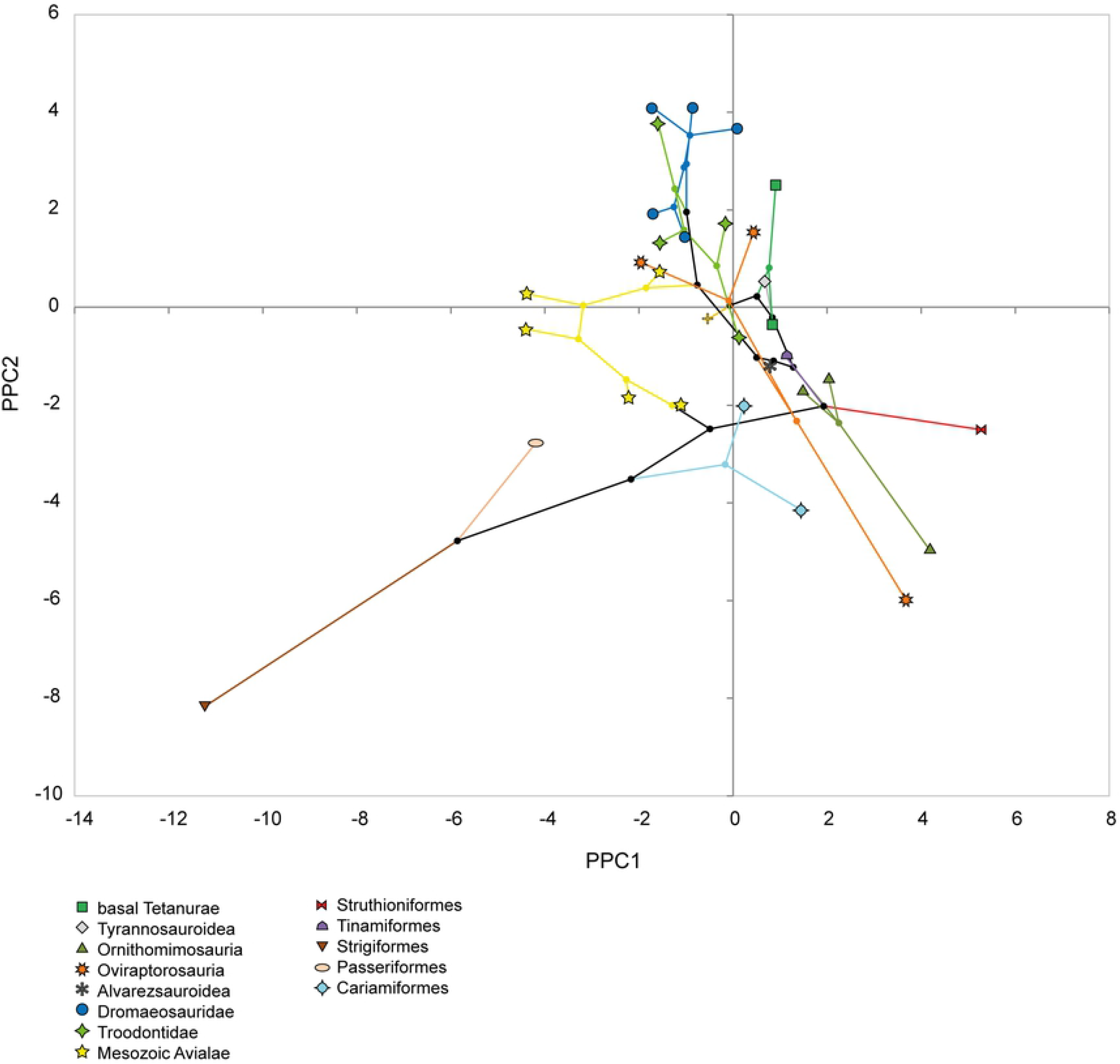
Phylomorphospace obtained from the phylogenetic PCA based on the lengths of the pedal phalanges (PPC1 vs PPC2).

**Fig 6.**
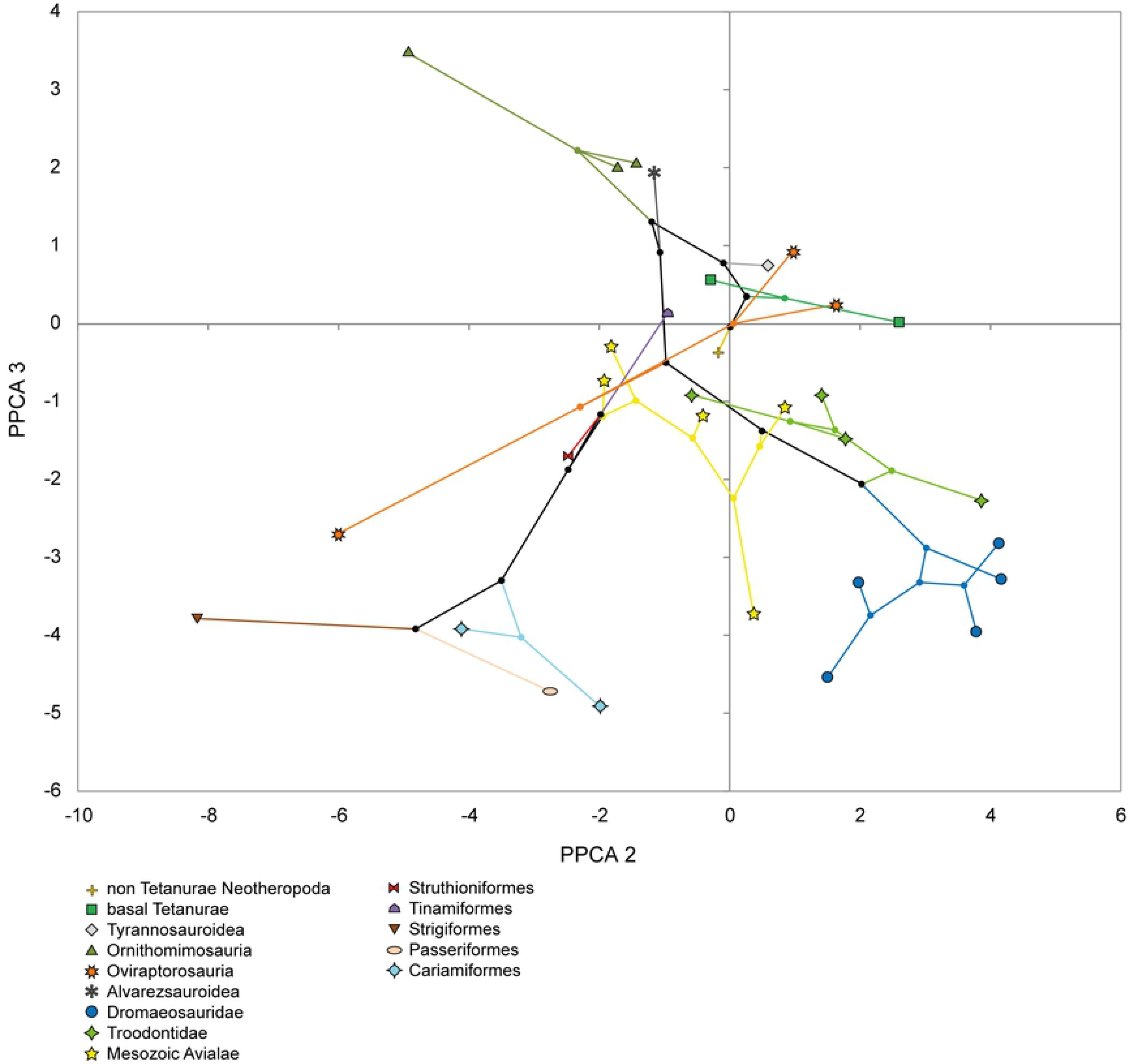
Phylomorphospace obtained from the phylogenetic PCA based on the lengths of the pedal phalanges (PPC2 vs PPC3).

Among oviraptorosaurs it can be observed a wide distribution of taxa along PPC3 (Fig 6), whereas in the remaining clades the distribution of taxa is more limited along this component according phylogenetic relationships. Thus, the basal taxon *Avimimus* is remarkably separated from the more derived *Corythoraptor* and *Khaan* mostly throughout the PPC3 and also the PPC2. So, the basal and derived taxa are mainly divided by length differences between the digit IV (larger in derived taxa) and the digits II and III (larger in basal taxa), as can be observed in the PPC2 vs PPC3 axes.

Regarding dromaeosaurids, they show a convergence between basal and derived taxa, since *Buitreraptor* is located near the derived eudromaeosaurids *Deinonychus* and *Bambiraptor* (Figs 5 and 6). These three taxa have a comparatively elongated digit IV than *Microraptor* and *Sinornithosaurus*, which are more derived than *Buitreraptor* although more basal with respect to the eudromaeosaurids.

Among troodontids there is observed an evolutionary trend to shorten the digit IV and to increase the length of digit III and to a slight elongation of proximal phalanges, as shows the PPC1vsPPC2 graphic (Fig 5). Thus the basal taxon *Anchiornis* is convergently located near to *Deinonychus* and *Buitreraptor*, with a proportionally more elongated digit IV, whereas *Troodon* has the shortest digit IV and a comparatively longer digit III as can be observed in the PPC2 vs PPC3 graphic (Fig 6).

Mesozoic avialans show a similar evolutionary trend than troodontids, since the basal taxon *Archaeopteryx* has an elongated digit IV whereas in more derived taxa this digit decreases in length whereas the other digits lengthen, specifically the digit III (Figs 5 and 6). In turn there is a trend to an elongation of the distal non-ungual phalanges of digits II, III, and IV in more derived forms, as can be observed in the PPC1vs PPC2 axes.

The sample of extant birds included in this analysis is small, although a certain evolutionary trend can be observed. In general lines, there is an increase in length of the distal non-ungual phalanges and of the digit III as a whole. Thus, the more basal *Rhea* has long proximal phalanges and a digit III comparatively shorter, whereas the more derived *Turdus* and *Bubo* have markedly longer distal non-ungual phalanges and a particularly elongated digit III.

### Influence of size in the distribution of taxa along the morphospace

Following the result of the PGLS regressions, the axes that compose the morphospace analyzed for the phalange measures (i.e., PPC1, PPC2, and PPC3) are not significantly influenced by size (PPC1: F = 1.253, p-value = 0.2722; PPC2: F = 2.513, p-value = 0.1238; PPC3: F = 0.6881, p-value = 0.6881). Accordingly, the distribution of taxa along the axes does not follow a pattern controlled by size.

## Discussion

Previous authors enumerated the morphological features of animals traditionally considered as ‘cursorials’: relatively long limbs; hinge-like joints; distal limb segments proportionally elongated; the reduction, compression or loss of the ulna and fibula and of the lateral metapodials and phalanges; reduction or loss of distal muscular groups or proximal location of their scars; a limb motion restricted to the sagittal plane;acquisition of digitigrade or unguligrade stance; and metapodials interlocked, fused or reduced to a single element [13, 56–61]. From the perspective of the locomotor performance, animals known as cursorials have the capacity to move at greater velocities or for extensive distances with a low energetic cost [60–63]. However, Carrano [61] considered that a discrete categorization of the locomotor habits could not be appropriate and instead these habits would be evaluated along a multivariate continuum between two locomotor extremes, i.e., strictly graviportal and cursorial. Theropods can be generally considered as cursorial animals (or ‘subcursorial’, according to Coombs [13]), due they were bipeds, digitigrades and with long and parasagittally oriented hindlimbs [64], although different taxa would be dispersed along a continuum that includes different grades of cursoriality. The distribution in the morphospace obtained in the multivariate analyses performed, could reflect such ecomorphological diversity. Thus, those taxa with more elongated distal segments of the hindlimbs (i.e., tibia and metatarsus), a more slender and compressed metapodium, and reduced lateral pedal digits likely had a greater cursorial capacity [59, 61]. These taxa would locate closer to the ‘cursorial extreme’ of the multivariate continuum than taxa with shorter segments of the hindlimb, with a more robust metapodium, and lateral digits less reduced.

The elongation of the distal elements of the hindlimb (i.e., tibia and metatarsus) allows increasing of the stride length and speed of movements, which are related to a greater cursorial capacity [9, 61]. Garland and Janis [60] explained than the ratio between the lengths of metatarsus and femur (MT/F) was repeatedly considered by some authors as a predictor of locomotor performance in fossil forms. However, Garland and Janis [60] and other authors [65–67] warned that ratios between hindlimb bones are not good predictors of the type of locomotion, so limb proportions must be considered with caution. Thus, it is important take to account also qualitative aspects, such as the morphology of the metapodium, to make inferences about locomotor capacities.

The arctometatarsal and subarctometatarsal conditions could confer significant cursorial capabilities. Some authors [12, 15] have verified that theropod taxa with these conditions have distal elements of the hindlimb significantly more elongated than taxa with a plesiomorphic metapodium. Moreover, many authors have postulated biomechanic hypotheses about the performance of the arctometatarsal and subarctometatarsal foot, and how the interaction motions between metatarsals and transfer of forces along the metatarsus provide advantages during locomotion, which could represent benefits for the cursorial habit [12–17, 68].

Regarding morphology of pedal phalanges, in extant terrestrial birds with a cursorial locomotor mode and walking capacity (e.g., ratites such as ostriches, emus, *Pterocnemia*, and *Rhea*) the pre-ungual phalanges tend to shorten distally [9, 69–71]. Further, in these birds the foot is symmetrical since digit III is the more developed and the main weight bearer, with non-ginglymoid interphalangeal articular surfaces, whereas digits II and IV have a similar length to each other, are shorten than digit III and have more ginglymoid interphalangeal articular facets, indicating that they were under higher torsional efforts [9, 72–73]. Similar features are observed especially in Mesozoic theropod taxa considered with greater cursorial capabilities, much of them possessing long tibiae and metatarsi and an arctometatarsalian condition, such as ornithomimids, alvarezsaurids, caenagnathids, and *Avimimus* (e.g., [9, 13, 33, 38, 55, 74–83].

By contrast, extant birds with a foot with grasping capacities are characterized by an elongation of the distal pre-ungual phalanges of the digits, especially the penultimate phalanx [69–71, 84–85]. This can be observed either in perching and raptorial extant birds. Even, the elongation of distal phalanges is convergently observed in arboreal mammals which have grasping autopodia, such as the sloths ([85], and references herein).

### Interpretation of the PPCA analyses related with the locomotor habits of theropods

Taking into account the diverse factors and how they affect differentially the hindlimb elements, it is important to consider both analyses together (i.e., long bones and phalanges proportions) to make adequate inferences about the locomotor habits of theropods. For instance, *Avimimus* and *Sinornithoides* are very close to each other in the PPCA morphospace constructed from the long bones measures, and there no evident differences (Fig 1), while the PPCA based on phalanges lengths reveals clear dissimilarities between these taxa (Figs 3 and 4). The later analysis indicates that the cursorial capacities of *Avimimus* are greater than those of *Sinornithoides*, whose phalanges proportions are possibly more related to a grasping function.

Based on the results of the PPCA made from phalanges length, taxa such as *Avimimus*, *Cariama*, and *Rhea* are considered with greater cursorial capacities [86–88], which are associated with more elongated proximal phalanges and a long digit III (Figs 3, 4 and 7). Other taxa, such as ornithomimids (especially *Struthiomimus*) also have traits related to more cursorial capabilities, i.e., more elongated proximal phalanges, although their digit III is not as long as in the taxa above mentioned. Instead, *Bubo*, *Turdus*, and some Mesozoic avialans close to them had a foot with elongated distal phalanges which possibly had more grasping capacities. Concerning taxa such as *Gualicho*, *Allosaurus*, *Gorgosaurus*, *Corythoraptor*, and *Khaan* they have slightly more elongated proximal phalanges, so could have had certain cursorial capacities, also taking into account they have a digit IV almost as long as digit III.

**Fig 7.**
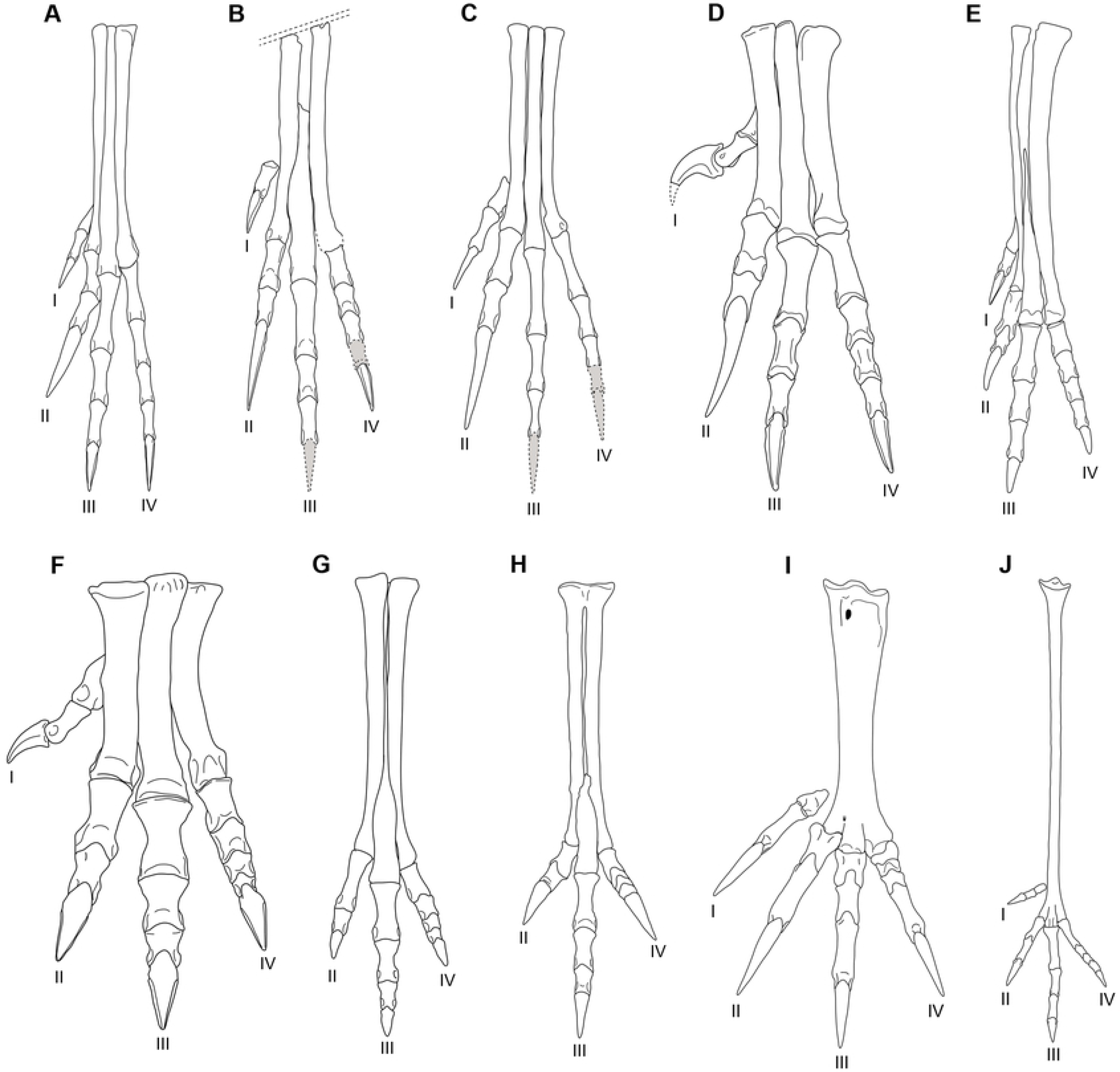
Comparison of the autopodium between several theropod taxa, including unenlagiines and some extant birds, in anterior view. (A) *Buitreraptor gonzalezorum* (based on MPCN-PV-598). (B) *Neuquenraptor argentinus* (based on the holotype, MCF-PVPH-77; phalanges III-4 and IV-4 lack in the original material). (C) *Rahonavis ostromi* (based on a cast of the holotype, FMNH PR 2830; phalanges III-4, IV-4 and IV-5 lack in the original material). (D) *Deinonychus antirrhopus*. (E) *Talos sampsoni*. (F) *Allosaurus gracilis*. (G) *Gallimimus bullatus*. (H) *Avimimus portentosus*. (I) *Bubo virginianus* (based on MACN 2056a). (J) *Cariama cristata* (based on MACN 23873). (A) is inverted from the original material to compare better to remain taxa. In (I) and (J) the first digit is showed disarticulated from its natural position (totally turned backwards) for better visualization. (D), (F) and (G) modified from Fowler et al. [9]; (E) based on Zanno et al. [151]; (H) based on Vickers Rich et al. [80].

The position of dromaeosaurids in the morphospace, including *Buitreraptor*, and other taxa, such as *Anchiornis*, is related to their long digit IV and elongated distal phalanges (Figs 3, 4 and 7). This feature could be related with their particular morphology where the digit II is markedly short and thus digits III and IV are the main structures of the foot support [6, 8, 89–94].

In the analysis based on long bones measurements (Fig 1) the PPC2 is less influenced by phylogeny and the distribution of taxa along this axis could show a clearer separation related with habits. TheMzTer on more positive values of PPC2 (*Allosaurus*, *Ceratosaurus*, *Beishanlong*, *Garudimimus*), which show short and robust metatarsi, can be considered with a minor cursorial capacity than those taxa tending to locate at less positive and negative values of PPC2 (*Dilong*, *Archaeornithomimus*, *Elaphrosaurus*, and *Herrerasaurus*), which show longer and slender metatarsi. In the case of modern birds also is observed the same general trend. Coincidentally, the taxa on negative values generally have comparatively smaller body sizes, except for *Megalosaurus*.

Along the PPC1 those taxa tending to positive or low negative scores can be considered with less cursorial capacities than those located at more negative scores. Thus, taxa such as *Linhenykus* and *Parvicursor*are interpreted with high cursorial abilities, in addition to having a markedly elongated and slender highly derived arctometatarsus [33, 78, 95] (Fig 8). Unfortunately, these two taxa have not preserved all the pedal phalanges and so they cannot be included in the analysis based on phalanges lengths. However, other alvarezsaurid considered in the analysis, i.e., *Kol ghuva* [96], shows pedal phalanges proportions that indicate cursorial capacities.

**Fig 8.**
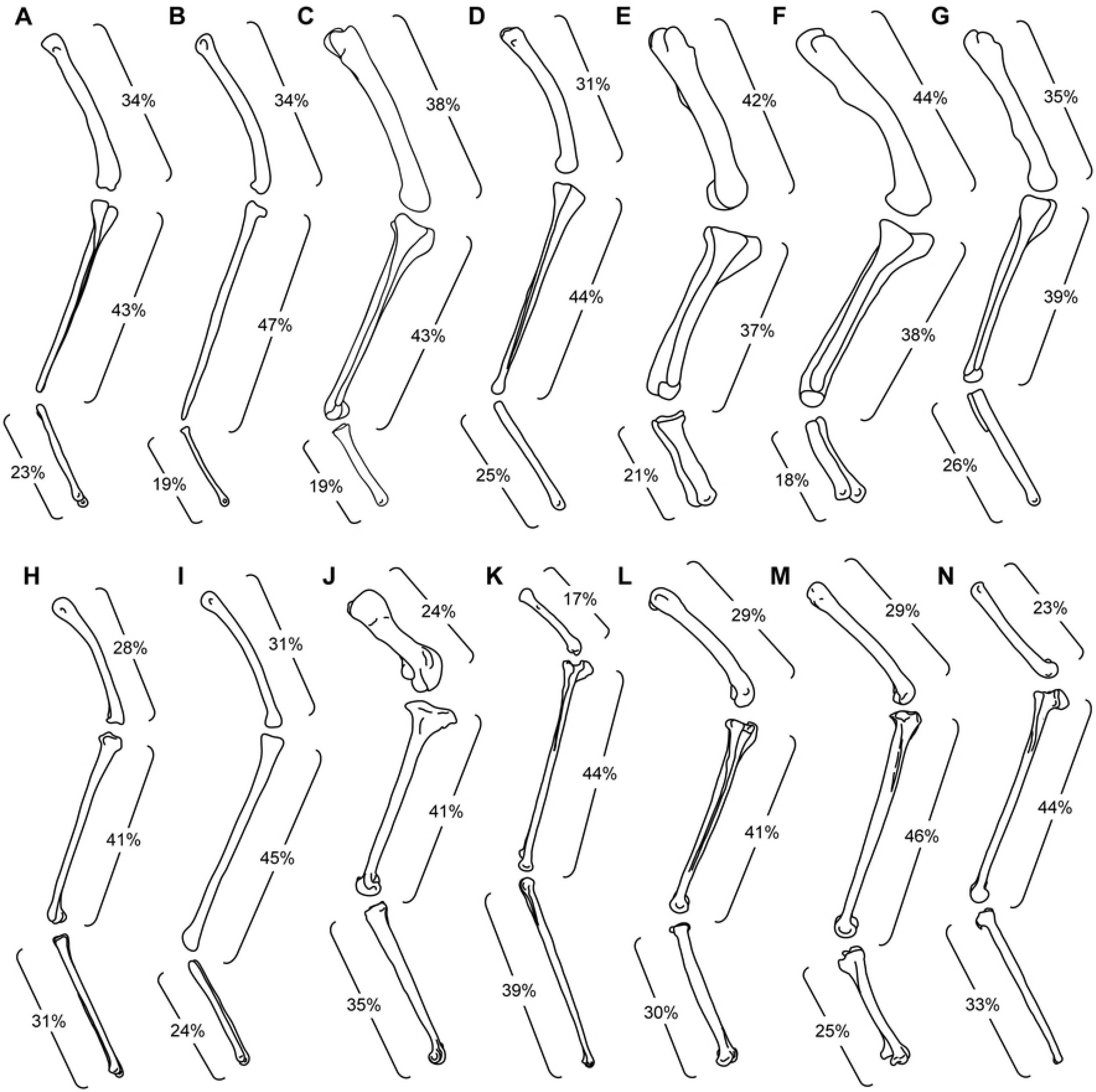
Comparison of hindlimb bones of different theropod taxa, including unenlagiines and extant birds, showing the proportional lengths of the femur, tibia and metatarsus. (A) *Buitreraptor gonzalezorum* (based on MPCN-PV-598). (B) *Rahonavis ostromi* (based on a cast of the holotype: FMNH PR 2830). (C) *Deinonychus antirrhopus*. (D) *Sinornithoides youngi*. (E) *Tyrannosaurus rex*. (F) *Allosaurus fragilis*. (G) *Struthiomimus altus*. (H) *Parvicursor remotus*. (I) *Archaeopteryx litographica*. (J) *Struthio camelus* (based on CFA-OR-1560). (K) *Cariama cristata* (based on MACN 23873). (L) *Geranoaetus melanoleucus* (based on MACN 2129). (M) *Bubo virginianus* (based on MACN 2056a). (N) *Furnarius rufus* (based on MACN 68647). Hindlimbs are not to scale. (C), (E) and (G) modified from Ostrom [43]; (D) based on Russell and Dong [152]; (F) modified from Gatesy and Middleton [65]; (H) based on Karhu and Rautian [78]; (I) based on Mayr et al. [153].

Our quantitative analyses, in addition to other features already described (subarctometatarsal configuration; [42, 97]) indicate that *Buitreraptor* can be considered with probable high cursorial capabilities. Other MzTer with probable similar locomotor capacities are the dromaeosaurids *Zhenyuanlong* and the troodontids *Sinovenator* and *Mei*, which were already described as possessing an arctometatarsal or subarctometatarsal condition [2, 42, 52, 97–100]. Further, these taxa present hindlimb and pes proportions similar to non-dromeosaurid theropods such as *Struthiomimus*, an ornithomimid probably markedly cursorial, as also indicate the PPCA based on phalanges lengths. Notwithstanding, *Buitreraptor* has phalanges proportions indicating grasping adaptations and related with a lesser cursorial performance. Unfortunately, phalanges lengths of *Sinovenator*, *Mei*, and *Zhenyuanlong* were difficult to obtain, because fragmentary preservation and incomplete information of the descriptions of the taxa, although in *Sinovenator* phalanges of digit III appear to shorten distally and the phalanx IV-4 is slightly longer than IV-3 [48].

### Functional implications of the dromaeosaurid hindlimb morphology and differences between unenlagiines and eudromaeosaurs

#### Long bones proportions, morphology of the metatarsus and motion range of digits

The main differences between the hindlimbs of unenlagiines and eudromaeosaurs are related with the relative length and form of the metatarsus, and the morphology of the phalanges of the digit II [4, 42, 101–102]. In unenlagiines the metatarsus is significantly elongated when is compared with the femur and tibia (except in *Rahonavis*), and it is slender because its lateromedial width (ML) is significantly lower than its total length (MtL) (except in *Rahonavis*) (Figs 7 and 8), whereas in eudromaeosaurs the metatarsus is remarkably shorter and the ratio ML/MtL is larger. Moreover, unenlagiines show a subarctometatarsal condition, whereas eudromaeosaurs have a metatarsus more similar to the plesiomorphic condition [6, 35, 44, 103–104]. These characters indicate that the metatarsus of eudromaeosaurs is overall more robust than that of unenlagiines.

The metatarsi of *Neuquenraptor* (MCF PVPH 77) and *Austroraptor* (MML 195) are incomplete, although their approximate length can be estimated, indicatingthey were very elongated with respect to the tibia and femur. Thus, these taxa possibly had length proportions of the hindlimb bones much similar to those of *Buitreraptor*. Moreover, *Neuquenraptor* and possibly *Austroraptor* (based on the specimen MML 220), also have a subarctometatarsal condition.

The long bone proportions of *Buitreraptor* are remarkably different with respect to those of eudromaeosaurs here analyzed, i.e., *Velociraptor*, *Deinonychus*, and *Bambiraptor* (Fig 8). Instead, *Buitreraptor* is more similar to other taxa with a relatively elongated metatarsus, either with an arctometatarsal, a subarctometatarsal, or non-subarctometatarsal condition, such as *Mahakala*, *Alnashetri*, *Zhongjianornis*, *Zhenyuanlong*, *Sinovenator*, and *Mei*. These taxa are similar in size or smaller than *Buitreraptor* [50, 52, 98–100, 105–108]. According to previous authors a similar size and hindlimb proportions would presumably indicate a similar locomotor mode [12, 15, 65]. Moreover, this resemblance in the locomotor mode can be also supported by the similar metatarsus morphology between some of these taxa.

*Rahonavis* departs from the general morphology of other unenlagiines, by its shorter tibia and a shorter and wider non-subarctometatarsal metatarsus (Figs 7 and 8) [109]. On the other hand, *Rahonavis* has hindlimb proportions more similar to those of unenlagiines than those of eudromaeosaurs, especially because it has a comparatively short femur and long tibia. Thus, *Rahonavis* can be considered as the less cursorial unenlagiine analyzed, although clearly more cursorial than eudromaeosaurs.

Additionally, differences in the distal articular surfaces of metatarsals between unenlagiines and eudromaeosaurs were also denoted by previous authors (e.g., [3, 9, 42]). In eudromaeosaurs the MT I, II and III have a well-developed ginglymoid distal articular surface [5, 6, 9, 44–45]. This could indicate that the first phalanges flexed and extended predominantly in a single plane [9]. Instead, in unenlagiines the ginglymoid distal facet of the MT II and III is less developed, so Ph. II-1 and III-1moved in a predominant vertical plane although probably with some degree of sideways movement. The distal surface of MT I of unenlagiines is ball-shaped, as in *Buitreraptor* (MPCA 238, [42, 97]) and *Rahonavis* (FMNH PR 2830) or it is slightly ginglymoid, as in *Neuquenraptor* [110–111].Thus, in *Neuquenraptor* the range of movement was probably more similar to that of digit I of eudromaeosaurs, whereas in *Buitreraptor* and *Rahonavis* digit I could have had a greater motion range. The more restricted motion of digits in eudromaeosaurs (which is emphasized by the more ginglymoid interphalangeal articulations in comparison with unenlagiines) could be more resistant to torsional stress and thus preventing disarticulation of the joints during manipulation of the prey with a greater grasping force [9]. In the case of the distal facet of MT IV it is generally more rounded in dromaeosaurids, which matches with the concave proximal articular facet of Ph.IV-1. This trait possibly indicates more freedom of movement for digit IV [9]. Thus, unenlagiines had the capacity to oppose pedal digits between them in a similar way to *Deinonychus* [9]. Digits I and IV probably had a wide range of motion, which would have allowed these digits converge during flexion, thus achieving a grip position.

### Morphology of pedal phalanges

The only unenlagiine with all the pedal phalanges preserved to date is *Buitreraptor*. Our results indicated that it is similar in phalanges proportions with respect to eudromaeosaurs analyzed, i.e., *Deinonychus* and *Bambiraptor*. The three taxa highlight by their markedly elongated digit IV, with a total length greater than that of digit III (Fig 7). In *Neuquenraptor* and *Rahonavis* it can be estimated that digit IV is shorter than digit III, as in *Sinornithosaurus* and *Microraptor*, because the sum of lengths of the other pre-ungual phalanges of digit IV is significantly lower than the total length of digit III and although Ph. IV-4 has been equal in length or slightly longer than Ph. IV-3 the complete digit IV would have been slightly shorter than digit III. By contrast in other MzTer included in the analysis such as derived troodontids, non-paravian coelurosaurs and basal tetanurans the digit III is the longest and the digit IV is significantly shorter, which are proportions related with probable more cursorial capacities [9, 72–73]. So, the length proportions of dromaeosaurids digits, including unenlagiines and especially *Buitreraptor*, seem to indicate a restriction to their cursorial habit.

Also, dromaeosaurids show a significant elongation of the distal pre-ungual phalanges, a feature related with grasping capacities (see cited literature above). Generally, in unenlagiines the length proportions of the distal phalanges of digit III are similar to those of eudromaeosaurs, although in the digit II the second phalanx is shorter than the first one (S1 Appendix), indicating slightly lower grasping capacities in unenlagiines. In others dromaeosaurids, such as *Microraptor*, Ph. III-3 is significantly shorter than III-2, a feature that also could indicate a decreasing of grasping capacities. Unfortunately, the lack of preserved elements prevents a more accurate analysis of the phalangeal proportions of *Neuquenraptor* and *Rahonavis*, although the available data and the apparently long distal phalanges of digit IV in *Neuquenraptor* indicate for this taxon more accentuated grasping capacities than other unenlagiines and resembling those of eudromaeosaurs (S1 Appendix).

In other groups of MzTer distal phalanges of digit IV generally maintain a similar length (S1 Appendix and S5 Text). By contrast, the length proportion of distal phalanges of digits II and III is more variable, due in some taxa these phalanges are shorter than the proximal ones (taxa considered as more cursorials) whereas in others taxa the distal phalanges are longer although they not surpass the length of the proximal ones (taxa with possible grasping abilities of the feet). In extant birds with a grasping foot, such as *Turdus* and *Bubo*, the distal phalanges are significantly long (S1 Appendix and Fig 7).

Many current birds with grasping capacities of the feet are ‘perchers’ and have arboreal habits, i.e., they are predominantly arboreal foragers [112]. An arboreal habit for some unenlagiines is difficult to envisage or impossible in taxa such as *Neuquenraptor*, *Unenlagia*, and *Austroraptor* because of their large sizes. Further, this habit is correlated in paravians with aerial locomotor capacities, although previous authors considered that aerodynamical features in large-sized dromaeosaurids were lost, as suggested by the scarce development or lack of papillae for feather attachment on the ulna [105]. In smaller taxa such as *Buitreraptor* and *Rahonavis* this lifestyle would have been more probable not only because of their smaller size but also because they have evidence of feathered forelimbs by preserved quill knobs (in *Rahonavis* [109]) and many osteological traits which suggest the capacity of flapping flight [4]. Also, it is possible that *Buitreraptor* and *Rahonavis* have been able to climb trees, especially considering the claw of pedal digit II as a potential tool for this purpose [7]. However, it is important to take into account the paleoenvironment in which they lived, since for example *Buitreraptor* was found in sedimentites that indicate a mainly aeolian environment and the existence of a large desert [113–116], where the trees were probably very scarce or nonexistent. So, the hindlimb morphology of *Buitreraptor*, mainly that of the metatarsus, is probably more related to a terrestrial habit than to an arboreal one.

Concerning qualitative aspects of the digit II of unenlagiines, it is modified as in eudromaeosaurs, although important differences are observed. First, in unenlagiines such as *Buitreraptor*, *Neuquenraptor*, and *Unenlagia paynemili* (MUCPv 1066), the distal articular surface of phalanx II-2 is less proximally extended. This feature restricts the extension of the ungual phalanx, as can be observed in an isolated articulated digit II of *Buitreraptor* (MPCA 478 [42]), in which the ungual seems to be totally extended (Fig 9). In *Deinonychus* and *Bambiraptor* this articular surface is more proximally extended (FAG personal observation of YPM 5205 and AMNH FR 30556), and thus the claw had the possibility of a greater extension (see [8]). Additionally, the phalanges of digit II of eudromaeosaurs are comparatively more robust than those of unenlagiines. This digit is the main implied in the predatory function, so a robust digit II in eudromaeosaurs could be advantageous to capture and subdue large prey. Moreover, eudromaeosaurs have a short Ph. II-1. Taking into account that the Ph. II-1 represent part of the out-lever of the flexor muscle of the digit (possibly the *M. flexor perforatus digiti II*, which probably was inserted onto the proximoventral zone of the phalanx as in extant birds [117–118]), the shortness of this phalanx could maximize mechanical advantage of the flexor and the grasping strength of digit II. Another difference is the more proximally extended proximoventral heel of phalanx II-2 of eudromaeosaurs, which possibly was an insertion point of flexor muscles [6].

**Fig 9.**
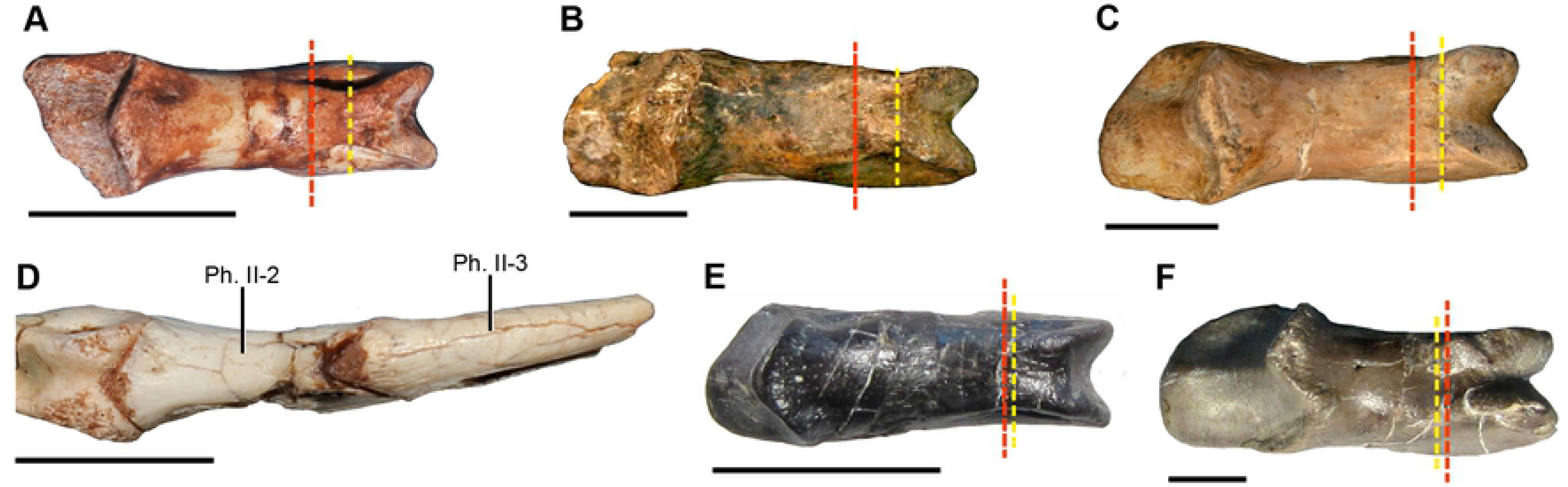
Comparison between pedal phalanges II-2 of unenlagiines and eudromaeosaurs, in dorsal view. The red dotted line indicates the posterior limit of the collateral ligament pit and the yellow dotted line indicates the posterior limit of the distal articular facet. (A) *Buitreraptor gonzalezorum* (MPCA 238). (B) *Neuquenraptor argentinus* (MCF PVPH 77). (C) *Unenlagia paynemili* (MUCPv 1066). (D) Articulated phalanges II-1, II-2 and II-3 of *Buitreraptor gonzalezorum* (the ungual phalanx is totally extended, so it is clear the proximal extent of the articular surface). (E) *Bambiraptor feinbergorum* (AMNH FR 30556). (F) *Deinonychus antirrhopus* (YPM 5205). Scale bars=1cm. (F) is courtesy of the Division of Vertebrate Paleontology; YPM VP.005205, Peabody Museum of Natural History, Yale University, New Haven, Connecticut, USA; peabody.yale.edu; photography by Federico A. Gianechini.

So, although in general traits the unenlagiines and eudromaeosaurs have phalanges of digit II with similar morphological characteristics, it is observed that these characters are more accentuated in the eudromaeosaurs, including a shorter phalanx II-1, a phalanx II-2 with a more proximally extended proximoventral heel, a shorter and more dorsoventrally constrained shaft, and a distal articular surface more extended proximally. This seems to indicate the presence of a digit II with the capacity of exert stronger predatory efforts in eudromaeosaurs, which could be an advantageous feature for subdue large preys. Conversely, the mentioned differences in the phalangeal morphology of unenlagiines could indicate weak predatory efforts, but the longer Ph. II-1 also suggests faster movements of digit II, what could be eventually useful for hunting small preys.

Regarding the degree of development and curvature of the claw of digit II it is difficult to evaluate differences between eudromaeosaurs and unenlagiines, mainly because most unenlagiines have not preserved a complete ungual. With the available data (S6 Table) we do not observe clear evidences indicating than eudromaeosaurs have a more developed and more curved ungual than unenlagiines.

Another possible difference between unenlagiines and eudromaeosaurs is respect to the location of digit I, which might have some implications in the grasping function. For instance, in *Deinonychus* the digit I is articulated to the middle zone of the diaphysis of the MT II [6, 9], suggesting it would have closed over the posterior face of the metatarsus during flexion. Moreover, previous authors proposed that in this taxon the metatarsus would have been positioned semi-horizontally while the animal was subject to its prey and thus helping to restrain it [9]. Among unenlagiines only one specimen of *Buitreraptor* (MPCN-PV-598) preserved a complete and articulated foot, in which the digit I seems to be located in the original position, articulated to the medial and distal surface of MT II [97]. This location could indicate that the metatarsus have been in a more vertical position during the submission of the prey, which would have been more effective for the digit I to participate in the gripping function.

### Morphological and functional correlates in extant raptorial birds and possible resemblances with dromaeosaurids

An interesting convergence is observed between extant raptorial birds and some eudromaeosaurs, in the morphospace of the long bone measurements. Both groups tend to positive PPC2 values (Fig 1), due they have longer femora and consequently shorter metatarsi. Moreover, raptorial birds converge specifically with *Deinonychus* and *Velociraptor* in the presence of wider metatarsi, as is reflected by their less negative values for PPC1 in both groups. Generally, in current raptorial birds a shorter and robust metatarsus is related with the ability of the foot to exert a greater grip force, whereas a longer metatarsus is related with a minor grip force although it has the capacity for rapid movement [9, 53, 66–67, 119–120]. In a general way, owls (Strigiformes) have the shortest and more robust metatarsus whereas falconids and especially accipitrids have a longer and slender metatarsus [53, 119–120]. Thus, owls have a greater grip capacity and strength, although these features also are related to other characters of the foot such as the presence of sesamoids, a specialized tendon locking mechanism and a facultative zygodactyl condition [53, 119–120]. Between the raptorial birds included in our analyses, *Milvago* and *Polyborus* (falconids of the subfamily Polyborinae) are characterized by relatively longer and slender tarsometatarsus when are compared with accipitrids (i.e., *Geranoaetus* and *Elanus*). This could indicate greater cursorial capacities, in agreement to what was expressed by previous authors [121].

Analogously, the short and robust metatarsus of eudromaeosaurs, such as *Velociraptor* and *Deinonychus*, could have allowed a great generation of grip force [6, 9]. By contrast, the elongated subarctometatarsus of unenlagiines could have had a greater capacity of rapid movement, like falconids and accipitrids, although it could have reduced grip strength [9].

Despite morphological and even functional features can be compared between these theropods and extant raptorial birds, it must be considered that these birds are predominantly aerial with a generally limited terrestrial locomotion (but see [121]). Many common features in the autopodium of raptorial birds can be interpreted as the result of a predominant influence of hunting and grasping specializations (e.g., elongation of distal non-ungual phalanges independently of the specific type of prey and the hunting method employed by them; [69–71, 85, 122], instead terrestrial locomotion. Conversely, dromaeosaurids, like most Cretaceous theropods, had a terrestrial locomotion, and it is expectable that both factor of selective pressures, i.e., predation and terrestrial locomotion, have a great influence in the hindlimb and autopodium. This is a main reason to explain the segregation in the morphospace between extant birds and dromaeosaurids, and also it might explain the presence of elongated distal phalanges in dromaeosaurids although not as strikingly long as those of extant raptorial birds (see also the study about the modular fashion of evolution of pedal phalanges proportions [85]). Thus, differences in hindlimb between eudromaeosaurs and unenlagiines can be considered mainly focusing in these, partially antagonist, specializations. The morphological design of the eudromaeosaurs autopodia indicates a more marked specialization to the predatory habit, whereas in unenlagiines a more marked cursorial specialization would have been occurred.

### Locomotor and predatory habits of *Buitreraptor* and other unenlagiines

Unenlagiines possibly had a better cursorial locomotor performance and the capacity to reach greater running velocities than eudromaeosaurs with shorter and more robust metatarsi. Of course, this does not mean that eudromaeosaurs did not have an effective locomotion and the ability to run fast, but that the morphological characters of the hindlimb of the unenlagiines would have given these animals greater and more efficient cursorial capacities. Possibly, eudromaeosaurs may have made sudden runs at high speeds, but for shorter periods of time or for short distances, while unenlagiines could maintain an accelerated pace for more time and/ or distance. Regarding the metatarsus of eudromaeosaurs it has a structure with functional capacities possibly more useful to predation than to cursorial locomotion. About the morphology of pedal phalanges the discrepancies observed between both groups, especially those of digit II, could be more directly related with different predatory habits.

Despite the mentioned dissimilarities between unenlagiines and eudromaeosaurs, it is remarkable that is the general structure of the metatarsus which shows a more drastic difference. The metapodium had a greater morphological plasticity along evolution of dromaeosaurids, since its structure differs significantly in unenlagiines and eudromaeosaurs (and in microraptorines, which also have a subarctometatarsal condition), in relation to the relative and different importance of the mechanical benefits associated both with predatory and locomotor functions in both clades. On the other hand, as was explained above, the length proportions of phalanges are not meaningfully dissimilar between these groups. This factor could be related to the phalanges are the main elements implied in predator functions, which exerted a greater selective pressure on their morphology, independently of the feeding strategy and locomotor habit. Nevertheless, some specific differences, such as the longer and slender phalanx II-1 and the greater freedom of movement of the remaining digits of unenlagiines, could allow them a fast and secure grip of small and agile/elusive prey that do not demand great efforts to be subdue.

Unenlagiines have similar modifications of the metapodium than microraptorines and probably they had a similar mode of moving on the ground, beyond the capacity of gliding postulated for some microraptorines [123–125]. It can be that these two groups of dromaeosaurids used digit II for predation, although the predatory habits, i.e., the hunting way and the type of prey, were not necessarily the same, also taking into account that microraptorines (at least *Microraptor* and *Sinornithosaurus*) have a phalanx II-1 shorter than II-2 (S1 Appendix), as in eudromaeosaurs. Moreover, some specimens of *Microraptor gui* indicate it fed mammals, enantiornithine birds, and fishes, which is evidence of diverse feeding habits and possibility of exploit different substrates such as ground, trees, and water, in that taxon [126–128].

Likely, unenlagiines preyed on rapid and elusive animals, although it is difficult to know more specifically the type of prey that they hunted, even without having direct evidences such as the gut content of *Microraptor* specimens. Nevertheless, it is possible to achieve an approximation of the feeding habits of unenlagiines, especially for the better represented taxa such as *Buitreraptor*. Regarding other unenlagiines the information is scarcer, so it is more difficult to infer if among them there were noticeable differences in the feeding modes and on the types of animals that preyed.

Considering the small size, slender proportions (especially those of metapodium), and the inferred cursorial capacities, *Buitreraptor* probably foraged on the ground searching small preys, such as invertebrates, reptiles or mammals, throughout large distances and probably employing high-speed pursuits in some cases. The fauna recorded from the fossiliferous area of La Buitrera, where *Buitreraptor* was discovered, also includes remains of small tetrapods such as snakes, sphenodonts, crocodyliforms, and mammals [129–133], which could have been potential dams. *Buitreraptor* would have employed its pes to subjugate and keep the prey immobile once it was reached. The fast movements and curved enlarged claw of digit II would have helped this function, and eventually causing serious injuries or even death of the prey.

Another reliable indicator of the type of diet and feeding strategy is the dental morphology. The teeth of *Buitreraptor* are numerous, tiny, lateromedially compressed, and devoid of denticles [134]. Instead, eudromaeosaurs are generally characterized by larger serrated teeth, such as *Dromaeosaurus*, *Deinonychus*, *Velociraptor*, *Saurornitholestes*, and *Tsaagan* [5–6, 35, 135–137], although many taxa have denticles only on the distal carina. These features would have allowed ingesting larger preys or tearing and cutting the flesh from them into smaller pieces. Feeding models were proposed for some taxa, such as *Deinonychus* [9], although they are difficult to apply to *Buitreraptor*because the size of their teeth and the lack of denticles. Due the latter feature and the absence of other flesh-tearing structures (e.g., the tomial tooth of extant raptorial birds) it is very likely that *Buitreraptor* has consumed whole small animals and that the teeth were mainly employed as a tool to hold the dams. Also is possible these teeth have been used to fragment small preys, to consume them in more than one swallow. In previous works it has been postulated that the dentition of *Buitreraptor* would indicate a piscivorous feeding mode [134]. Certainly, *Microraptor* also has small non-serrated teeth and there is evidence that it fed on fishes. However, this unique feature is not a reliable indicator of piscivory, since other morphological evidences must be taken into account. Moreover, *Microraptor* included in its diet other animals in addition to fish, as mentioned above. *Buitreraptor* also is characterized by having long forelimbs and hands [4, 97], so it could also have used them to handle the prey once it was captured and subjugated with its feet.

Among extant long-legged predominantly terrestrial birds that forage on the ground and hunt small preys are included the seriemas (Cariamiformes) and the secretary bird (Falconiformes). The secretary bird kicks and stamp on the prey until it is wounded or incapacitated and then takes it with its beak [138–140]. On the other hand, the red-legged seriema (*Cariama cristata*) takes the prey with the beak and hits it on the ground with sudden movements of the head until it is injured [141]. An interesting trait of this seriema is it has a markedly curved ungual phalanx in the second digit ([142–143]; FAG personal observation of MACN 23873). Some authors proposed this bird use this claw to hold the prey against the ground, although others do not agree ([86], and references therein). The extinct phorusrhacids were terrestrial generally flightless carnivorous birds which also are characterized by having a markedly developed and curved ungual of the second digit [143–145]. Some authors have proposed that this claw could be used as a means of apprehension of the prey on the substrate, then using the beak to tear it apart [143]. *Buitreraptor* could be used its pedal claw in a similar way than that proposed for seriemas and phorusrhacids, although there are no direct evidences.

Regarding other unenlagiines such as *Austroraptor* it could be proposed a similar strategy of hunting and subjection of the dam than that of *Buitreraptor*. Although *Austroraptor* is significantly larger (estimated total length: 5 m) it has numerous and small teeth in comparison with the size of the skull, and also they lack denticles [134, 146–148]. However, the teeth of *Austroraptor* are conical, so they probably were more resistant and could have employed to retain and dismember large prey. Due to *Austroraptor* probably had similar length proportions of the hindlimb bones than *Buitreraptor* and a subarctometatarsal condition it could have had good cursorial capacities. By other side, *Austroraptor* has strikingly shorter arms than other unenlagiines, so it would not have used them to manipulate the prey, or at least not in the same way that *Buitreraptor*.

*Rahonavis* probably had less cursorial capacities due its hindlimb morphology, although it has a relatively long tibia, so fast chases of preys cannot be ruled out as a hunting strategy used by this taxon. Moreover, *Rahonavis* has a digit II similar to that of other unenlagiines, so it probably had similar functional capacities. However, the distal phalanges are shorter than in other unenlagiines, so it probably had slightly lesser gripping abilities. Unfortunately cranial remains and teeth of *Rahonavis* are unknown, so it is more difficult to speculate about the type and size of animals that it could have been preyed upon. Surely it fed on small preys, although is not possible to know if it was able to tear flesh from larger preys. Cranial remains neither were preserved in *Neuquenraptor*, although the features of its hindlimb indicate that velocity probably was important to obtain its preys. Regarding others unenlagiines, such as *Unenlagia comahuensis* and *Unenlagia paynemili*, they have not preserved cranial bones although have preserved scarce hindlimb remains, especially phalanges of digit II, which are much similar to those of the other unenlagiines [101–102, 149–150]. So, mainly due to the lack of skull and metatarsus remains and most of pedal phalanges it is more difficult to infer locomotor and predatory habits of these two species.

## Conclusions

Morphological differences in the hindlimb between unenlagiines and eudromaeosaurs reflect differences both in locomotor and predatory habits. In unenlagiines the presence of a long tibia and a long, slender and subarctometatarsal metatarsus suggest greater cursorial capacities with respect to eudromaeosaurs, which have a shorter, wider and non-subarctometatarsal metatarsus. Regarding pedal digits the two groups of dromaeosaurids have similar length proportions and based on the elongation of the distal phalanges they probably have the capacity of grasping. However, morphological features of eudromaeosaurs, i.e., a more robust metatarsus; distal articular surfaces of metatarsal I, II, and III, and interphalangeal articular surfaces markedly ginglymoid; and a shorter phalanx II-1, indicate that these dromaeosaurids possibly exerted more grip strength than unenlagiines. By contrast, proportions and slenderness of unenlagiines would not have allowed them to perform high grasping forces but instead they may have been able to make faster movements with both the metatarsal and the digit II. Moreover, this morphofunctional difference is analogously observed in extant raptorial birds, since in the latter those taxa with the shortest metatarsus, such as owls, have the ability to produce the greatest grip force, whereas those taxa with longer metatarsi, such as polyborine falconiforms, generate a lesser grip force although can effect faster movements with the pes.

Despite the presence of morphological differences of pedal phalanges between unenlagiines and eudromaeosaurs, these discrepancies are not as drastic as those observed between the metatarsus of both dromaeosaurids groups. This, together with the similar length proportions of pedal phalanges seem to indicate that the morphology of these pedal elements varied scarcely along dromaeosaurid evolution, a factor probably related with a greater selective pressure exerted by the predatory function.

Among unenlagiines, *Buitreraptor gonzalezorum*, with its small size, high cursorial capacities, a long metatarsus and phalanx II-1, more mobile phalanges, and tiny teeth, probably was a terrestrial predator that preyed on small elusive animals, such as arthropods, lizards, mammals, etc., trough rapid movements of its pes. *Rahonavis ostromi* also was a small-sized unenlagiine, although its morphology seems to indicate it had lesser cursorial abilities. Probably, its small body size and potential capacity of climbing could capacitate it to an arboreal habit. Other unenlagiines, such as the large-sized *Austroraptor cabazai* and the medium-sized *Neuquenraptor argentinus* probably preyed on larger animals, also making use of its high cursorial faculties. Regarding other taxa, such as *Unenlagia comahuensis* and *Unenlagia paynemili*, are more fragmentary and so is much difficult to infer a locomotor and predatory habit.

Along dromaeosaurid evolution the different lineages seem to have diverged in varied lifestyles, as is documented by unenlagiines, microraptorines, eudromaeosaurs, and recently by halszkaraptorines [11]. Future studies, such as reconstructions of the muscular system, will be necessary to analyze the hindlimb as an osteo-muscular integrated complex and how it would have been involved both in locomotion and depredation in dromaeosaurids. These paleobiological aspects will help us to have a better comprehension of the dromaeosaurid evolutionary story and about the role of these theropods within the ecosystems in which they lived.

## Acknowledgments

We thank to the curators of different institutions, who kindly allow us access to the materials of the collections under their care: Pablo Tubaro, Darío Lijtmaer, and Yolanda Davies, from the División de Ornitología, MACN, Buenos Aires, Argentina; Sergio Bogan, from Fundación de Historia Natural “Félix de Azara”, Universidad Maimónides, Buenos Aires, Argentina; Daniel Cabaza, from the MML, Río Negro, Argentina; Carlos Muñoz, from the MPCA, Cipolletti, Río Negro, Argentina; Pablo Chafrat, from the MPCN, General Roca, Río Negro, Argentina; Daniel Brinkman, from the YPM, Connecticut, USA; and Carl Mehling, from the AMNH, New York, USA.

Thank you to Washington Jones for sharing kindly information and bibliography about phorusrhacids and seriemas, and Nestor Toledo for share his ideas about functional aspects of tetrapod hindlimb.

## Supporting information

**S1 Appendix. Database including measurements of taxa used for the phylogenetic principal component analyses.** Measurements include those of long bones of the hindlimb and pedal phalanges lengths.

**S2 Fig. Example figure showing the methodology for measuring curvature angles of ungual pedal phalanges.**

**S3Table. Results of the phylogenetic principal component analysis based on long bones measurements.**

**S4 Table. Results of the phylogenetic principal component analysis based on phalanges lengths.**

**S5Text. Supplementary discussion.**

**S6Table. Curvature angles of pedal ungual phalanges of dromaeosaurids.**

